# Isolating Small Extracellular Vesicles from Small Volumes of Blood Plasma using size exclusion chromatography and density gradient ultracentrifugation: A Comparative Study

**DOI:** 10.1101/2023.10.30.564707

**Authors:** Fang Kong, Megha Upadya, Andrew See Wong Weng, Rinkoo Dalan, Ming Dao

**Affiliations:** School of Biological Sciences, Nanyang Technological University, SINGAPORE; Facility for Analysis, Characterisation, Testing and Simulation, Nanyang Technological University, SINGAPORE; Lee Kong Chian School of Medicine, Nanyang Technological University, SINGAPORE; Department of Material Science and Engineering, Massachusetts Institute of Technology, USA

## Abstract

Small extracellular vesicles (sEVs) are heterogeneous biological vesicles released by cells under both physiological and pathological conditions. Due to their potential as valuable diagnostic and prognostic biomarkers in human blood, there is a pressing need to develop effective methods for isolating high-purity sEVs from the complex milieu of blood plasma, which contains abundant plasma proteins and lipoproteins. Size exclusion chromatography (SEC) and density gradient ultracentrifugation (DGUC) are two commonly employed isolation techniques that have shown promise in addressing this challenge. In this study, we aimed to determine the optimal combination and sequence of SEC and DGUC for isolating sEVs from small plasma volumes, in order to enhance both the efficiency and purity of the resulting isolates. To achieve this, we compared sEV isolation using two combinations: SEC-DGUC and DGUC-SEC, from unit volumes of 500 *μl* plasma. Both protocols successfully isolated high-purity sEVs; however, the SEC-DGUC combination yielded higher sEV protein and RNA content. We further characterized the isolated sEVs obtained from the SEC-DGUC protocol using flow cytometry and mass spectrometry to assess their quality and purity. In conclusion, the optimized SEC-DGUC protocol is efficient, highly reproducible, and well-suited for isolating high-purity sEVs from small blood volumes.

## 1. Introduction

In recent years, extracellular vesicles (EVs) have garnered significant scientific attention due to their potential as biomarkers and applications in targeted drug delivery [1]. These vesicles are secreted by all cell types and have been detected in various types of bodily fluids such as blood, urine, saliva, and feces [2]. Comprising a phospholipid bilayer with a composition similar to the cell of origin, EVs carry a diverse range of molecules, including proteins, nucleic acids, and lipids [3]. They play crucial roles in both normal physiological processes and pathological conditions.

EVs can be classified into three main subgroups based on their biogenesis: exosomes, microvesicles (MVs), and apoptotic bodies. Due to the challenges in distinguishing between exosomes and MVs smaller than 150 nm, which share similar size, density, and protein markers, they are collectively referred to as small extracellular vesicles (sEVs) [4]. These sEVs are of particular interest in the field of biomarker research and targeted drug delivery, given their ubiquitous presence and functional relevance in various biological processes.

Blood plasma sEVs provide valuable information for diagnosis, prognosis, and homeostasis [5]. However, isolation of sEVs from plasma is hindered by two major constituents: plasma proteins and lipoproteins [6–8]. Contamination of these constituents in sEV isolates can affect the understanding of the role of sEVs as carriers of genetic information and protein antigens in human physiology and pathology. Numerous techniques, including differential ultracentrifugation (dUC), size exclusion chromatography (SEC), ultrafiltration, immunoaffinity isolation, and microfluidics [9], are used to isolate sEVs from plasma. SEC [10] and density gradient ultracentrifugation (DGUC) are particularly popular due to their unique advantages in isolating sEVs based on size and buoyant density, respectively. In many occasions involving human blood, such as routine blood tests, only a small volume of blood (1∼10 *ml*) is collected, which yields a limited amount of plasma (500 *µl* to 5 *ml*). To obtain high-purity sEVs with a reasonable yield from this small volume of plasma, it is essential to establish a repeatable and reliable isolation protocol. Additionally, the isolation method should be simple, practical, and involve commonly available equipment to ensure ease of use. To address this challenge, we explored the combination of SEC and DGUC, which has been shown to lead to high-purity sEV isolates by effectively removing both lipoproteins and plasma proteins [11–15]. However, the ideal sequence for these methods in isolating sEVs from small volumes of plasma remains unclear. In this study, we compared the sequences of SEC-DGUC and DGUC-SEC to determine the better option in terms of purity and yield. Our study reinforces previous findings [13,16] on the efficacy of size exclusion chromatography (SEC) in eliminating plasma proteins and high-density lipoproteins (HDLs), as well as the utility of density gradient ultracentrifugation (DGUC) in distinguishing small extracellular vesicles (sEVs) from lipoproteins.

We examined the performance of two protocols, SEC-DGUC and DGUC-SEC, in terms of sEV purity and yield. Both methods produced high-purity sEVs, but the SEC-DGUC protocol outperformed DGUC-SEC in terms of sEV protein and RNA yield.

In order to enhance the efficiency of the DGUC process, we optimized it using a tailored density gradient in a 1.5 *ml* tube format and a fixed-angle rotor. This adjustment significantly reduced the ultracentrifugation time to just 2 hours. Our optimized approach enabled the harvesting of sEVs across an extensive density range (> 1.08 *g/ml*), achieving high-purity isolation in a short span of 3 hours. This study thus offers an efficient protocol for sEV isolation that combines optimal yield and high purity. Subsequently, we further characterized the isolated sEVs using flow cytometry and mass spectrometry techniques. These additional investigations provided further confirmation of the sEVs’ purity and structural integrity.

In conclusion, our study highlights the superiority of the SEC-DGUC protocol over DGUC-SEC in terms of purity and yield when isolating sEVs from as little as 500 *µl* of plasma. This optimized protocol is well-suited for clinical applications requiring high-purity sEVs from small blood volumes and is easily adaptable for various research and clinical settings due to its simplicity, practicality, and use of commonly available equipment.

## 2. Result

### 2.1 SEC isolated particles from plasma with 1% sEV content

We first assessed the efficacy of SEC in isolating particles, particularly sEVs, from plasma (Figure 1A). A volume of 500 *µl* plasma was loaded onto a SEC column with a maximum loading capacity of 2 *ml*. We collected and pooled four fractions, fraction 7-10 (2 *ml* total volume), as the “particle fraction” (PF) for each SEC run. This procedure is consistent with the particle concentration profiles (Supplementary Figure 1A) and follows the SEC column manual’s guidelines. NTA measurement of these PFs, obtained from 10 blood plasma samples, demonstrated particle concentrations ranging from 9.6×10^8^ to 5.5×10^11^ *ml^-1^*.

**Figure 1:**
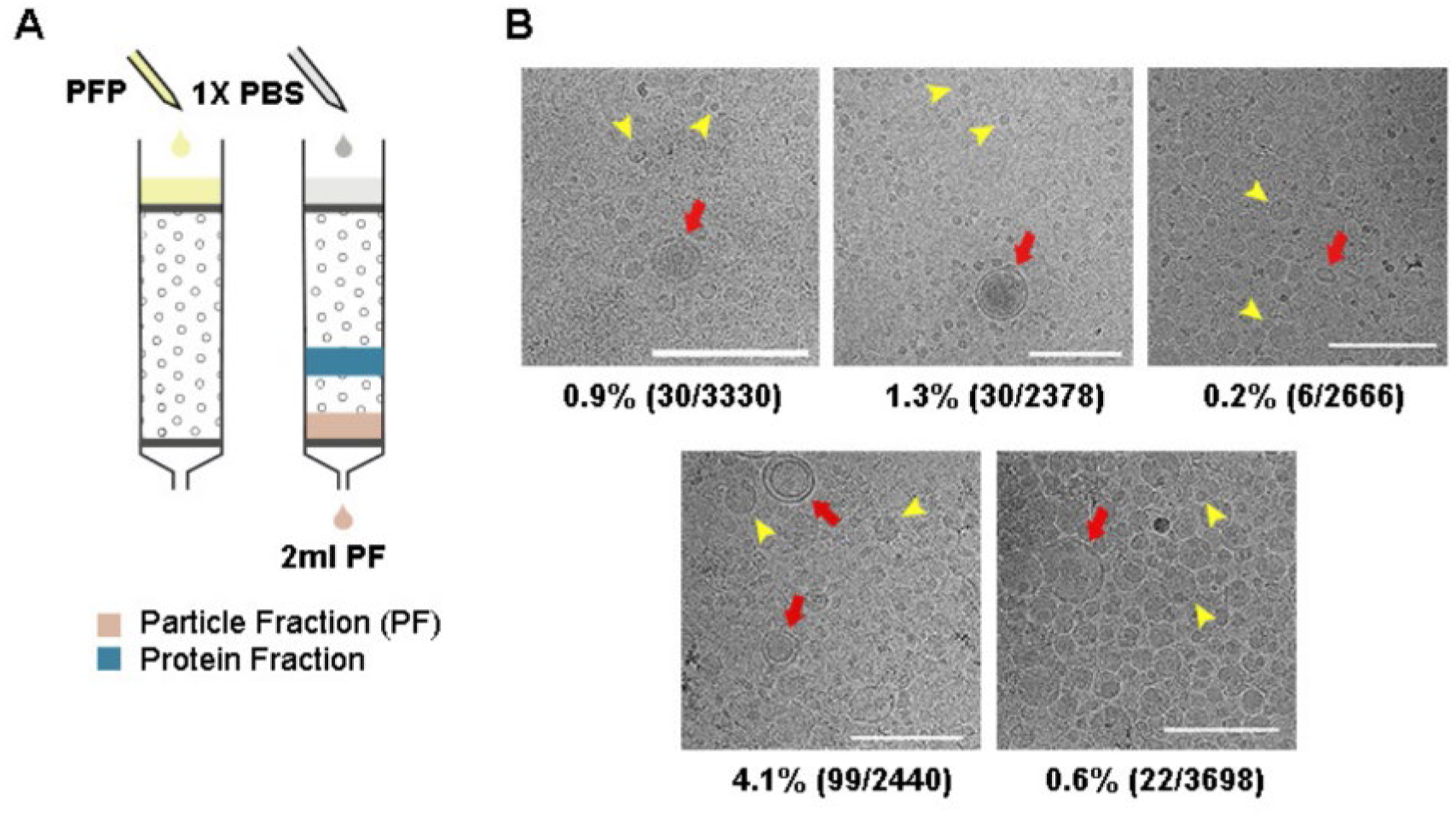
SEC was effective in removing plasma proteins and HDL but not low-density lipoproteins. (A) 500 *μl* of plasma was loaded on a SEC column and PFs were collected. (B) Cryo-EM images of PFs obtained from the plasma of 5 healthy individuals. Vesicles could be easily identified and the images revealed a clean background suggesting minimum protein contamination. Moreover, HDL was sparse in the Cryo-EM images, indicating that they were also largely removed. Furthermore, sEVs, having clearly defined bilayer (red arrows), were easily distinguishable from lipoproteins (representative particles marked by yellow arrows). The ratio of counted sEVs to all vesicles are shown under each image. All scale bars represent 200 *nm*.

Cryo-EM images of PFs isolated from the plasma of 5 healthy individuals were used to analyse the content of the PFs (Figure 1B). We observed distinct sEVs characterized by bilayer membranes, although they were outnumbered by a large number of lipoproteins (Figure 1B and Supplementary Figure 1C). The average proportion of sEVs in the PFs was 1%. Notably, HDLs (7-13 nm) [17] were observed quite sparingly in the cryo-images of PFs, suggesting a substantially lower presence compared to the other particle populations. On the other hand, the size of the lipoproteins in PF mainly comprised two populations centred at 20-25 nm and 40-50 nm, corresponding to the sizes of LDL and IDL/VLDL, respectively (Supplementary Figure 1F).

Our data demonstrated that SEC effectively removed plasma proteins and HDLs. However, sEVs constituted only 1% of the particles in the PFs, with the remaining 99% consisting of a mixture of LDL, IDL/VLDL lipoproteins.

### 2.2 DGUC density gradient design inside a 1.5 *ml* Eppendorf tube

Addressing the challenges associated with handling small volumes of plasma in DGUC requires an optimized approach, ideally involving the use of a smaller tube format, such as a 1.5 *ml* Eppendorf tube. For such optimization, it is critical to accurately identify the density zones of sEVs and lipoproteins, including their overlapping regions, to ensure effective separation.

To fulfil this initial objective, a DGUC experiment was conducted using a swing-bucket in a conventional 12 *ml* tube. The PF from SEC of 6 *ml* plasma served as the starting material, ensuring ample substance for subsequent TEM analysis (Supplementary Figure 2, Materials and methods). After the DGUC, the 12 *ml* tube was fractionated into 42 distinct fractions, each of which underwent NTA, TEM, and density measurements. This enabled correlation of the density profile with particle concentrations and TEM images, fostering an understanding of how sEVs separate from lipoproteins within the established density gradient (Supplementary Figure 2). The analysis delineated three discrete density zones. The first zone, with densities less than 1.05 *g/ml*, was characterized by high particle counts and a predominance of lipoproteins, yet devoid of sEVs. The second, a separation zone, spanning from 1.05 *g/ml* to 1.08 *g/ml*, displayed a reduction in particle numbers, emergence of sEVs, and a continued dominance of lipoproteins. The final zone, extending beyond 1.08 *g/ml* and up to 1.2 *g/ml*, marked as the sEV zone, exhibited a further drop in particle counts, a sharp decrease in lipoproteins, and a marked increase in the presence of sEVs.

Building on the findings from the 12 *ml* tube format experiment, we aimed to establish a density gradient within a more compact, 1.5 *ml* tube. This was accomplished by layering an 800 *µl* 10% iodixanol solution over a 20 *µl* 50% iodixanol cushion (Figure 2A). Subsequently, the assembled density gradient was calibrated with an additional 500 *µl* of PBS followed by DGUC, utilizing a fixed-angle rotor for durations of 2, 6, and 16 hours (see Materials and methods).

**Figure 2:**
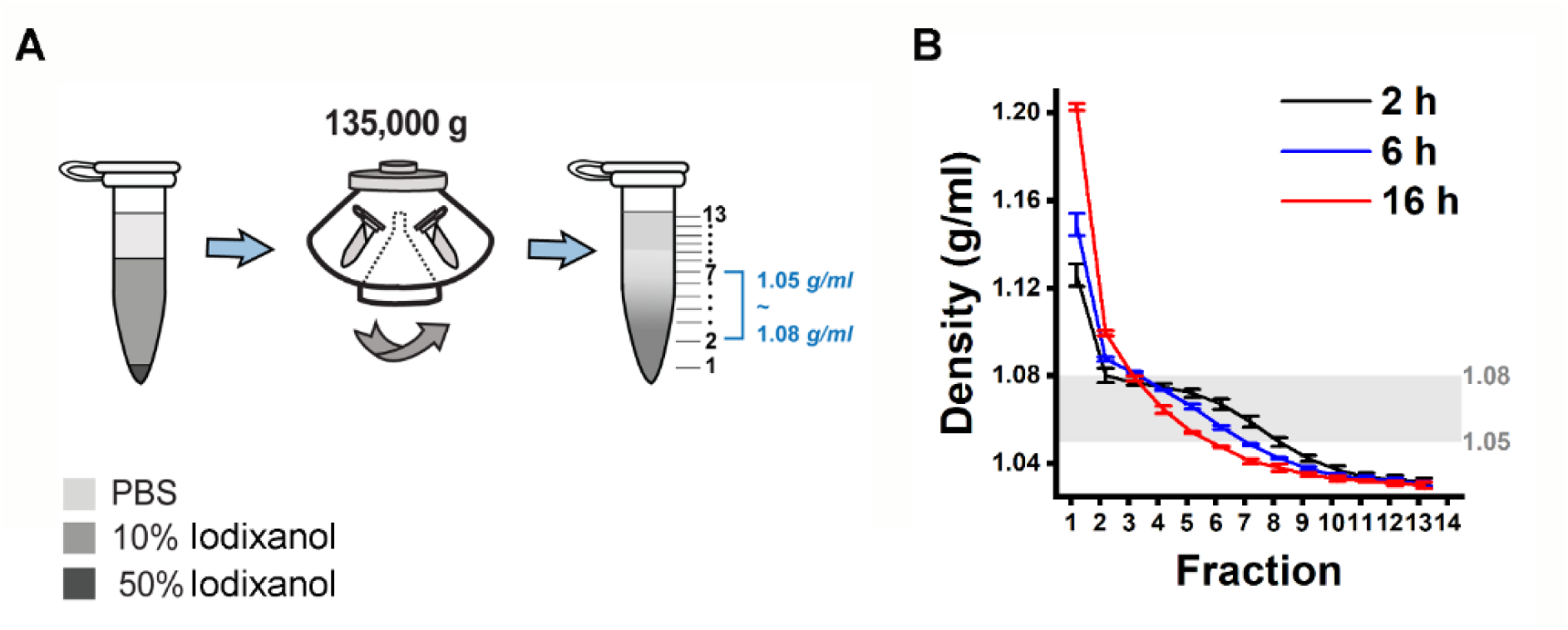
Design of a density gradient setup in a small volume format. (A) 500 *μl* PBS was overlaid on 800 *μl* 10% iodixanol solution and 20 *μl* 50% iodixanol cushion. The 1.5 *ml* tubes were subjected to ultracentrifugation using a fixed-angle rotor at an average speed of 135,000 *g* for 2, 6 and 16 *h* at 4 *°C* to establish a density gradient profile. (B) Density gradient profiles along the 1.5 *ml* tubes after ultracentrifugation. The 2 *h* spinning time gave a density profile with the largest separation zone of 1.05∼1.08 *g/ml* and smallest sEV zone of >1.08 *g/ml*. The data shows the average ± standard deviation of 7 repeats for 2 *h*, 3 repeats each for 6 and 16 *h*.

Following the small-tube format DGUC, the tube was fractionated into 13 parts of 100 *µl* each, with the first fraction at 120 *µl* to include the 20 *µl* 50% iodixanol cushion (Figure 2A). The density profiles displayed notable consistency across different runs at the tested time intervals of 2, 6, and 16 hours (Figure 2B). As the ultracentrifugation duration increased, the profiles evolved from a stepwise to a more continuous pattern. Within the 1.5 *ml* tube, the potential separation zone (1.05∼1.08 *g/ml*) slimmed down from a region of 700 *µl* to 500 *µl* and 300 *µl* as the ultracentrifugation period extended from 2 to 6 and 16 hours, respectively. Concurrently, the potential sEV zone (>1.08 *g/ml*) expanded from 120 *µl* during a 2-hour ultracentrifugation to 220 *µl* for both 6 and 16 hours. These findings indicated that a 2-hour ultracentrifugation period resulted in the broadest separation zone and the smallest sEV zone. Given the insights gained from these experiments, it appears that optimizing the DGUC process for a smaller, 1.5 *ml* tube format and initially setting the ultracentrifugation period at 2 hours could be effective for isolating sEVs.

### 2.3 DGUC isolated sEVs with significant protein contamination

Building upon these observations, the effectiveness of the 1.5 *ml* tube format DGUC was evaluated for direct sEV isolation from plasma. A 500 *µl* plasma sample was subjected to the 2-hour DGUC protocol, with the tube subsequently divided into 13 fractions. Each fraction’s particle and protein concentrations were subsequently measured (Figure 3). The high-density bottom fraction (>1.08 *g/ml*), which contained sEVs, was denoted as plasma-DGUC-1. The particle and protein concentration profiles revealed that the plasma-DGUC-1 fraction successfully eliminated the majority of proteins and a significant number of particles (Figure 3). However, despite the substantial protein removal, the protein concentration in plasma-DGUC-1 remained relatively high at 7.7 *mg/ml*. This indicates that while DGUC effectively separates low-density particles from those of high density, a considerable amount of protein still remains within the sEV density zone.

**Figure 3:**
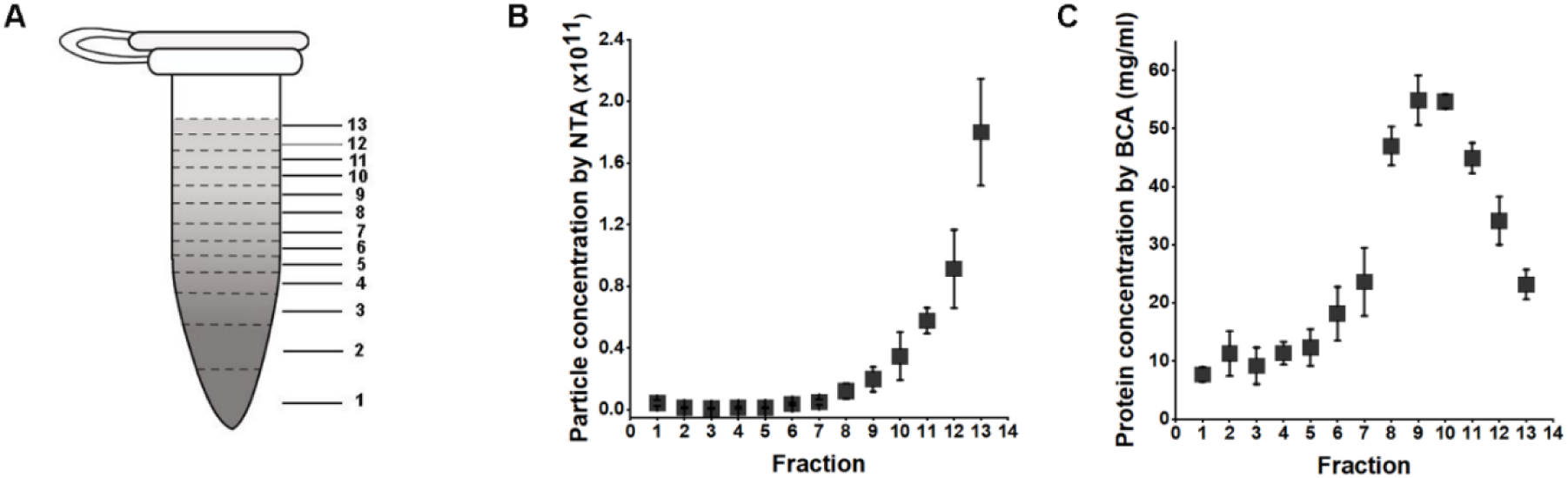
Particle and protein concentrations along the 1.5 *ml* tube after applying DGUC to plasma. (A) The 1.5 *ml* tube was fractionated into 13 fractions. (B) Particle concentration ± std measured by NTA of different fractions (n=3). (C) Protein concentrations ± std measured by BCA of different fractions (n=3).

### 2.4 SEC-DGUC protocol isolated high-purity sEVs

Recognizing the unique advantages and drawbacks of SEC and DGUC, it became clear that a strategic integration of these methods could potentially enhance the isolation of high-purity sEVs from blood plasma. With this perspective, we adopted a combined SEC-DGUC method. Specifically, we subjected PFs, obtained from SEC of a 500 *µl* plasma sample, to the small-tube format DGUC protocol (Figure 4A). The tube contents were systematically divided into 13 fractions, each of which was subjected to NTA, TEM, and density measurements. As expected, fraction 1, comprising the bottom 120 *µl*, showed a high density (1.10 *g/ml*) and was populated with high-purity sEVs (Figure 5). Fractions 2 to 7, within the 1.05∼1.08 *g/ml* density zone, displayed sparse sEV presence amidst lipoproteins, while the remaining fractions (8-13) were dominated by lipoproteins (Figure 5 and Supplementary Figure 3). Distinct NTA size histograms for fraction 1 further highlighted its divergence from the remaining fractions, which were overwhelmed by lipoproteins (Supplementary Figure 4B). As a result, fraction 1 was identified as the final sEV isolate derived from the plasma sample using the SEC-DGUC protocol.

**Figure 4:**
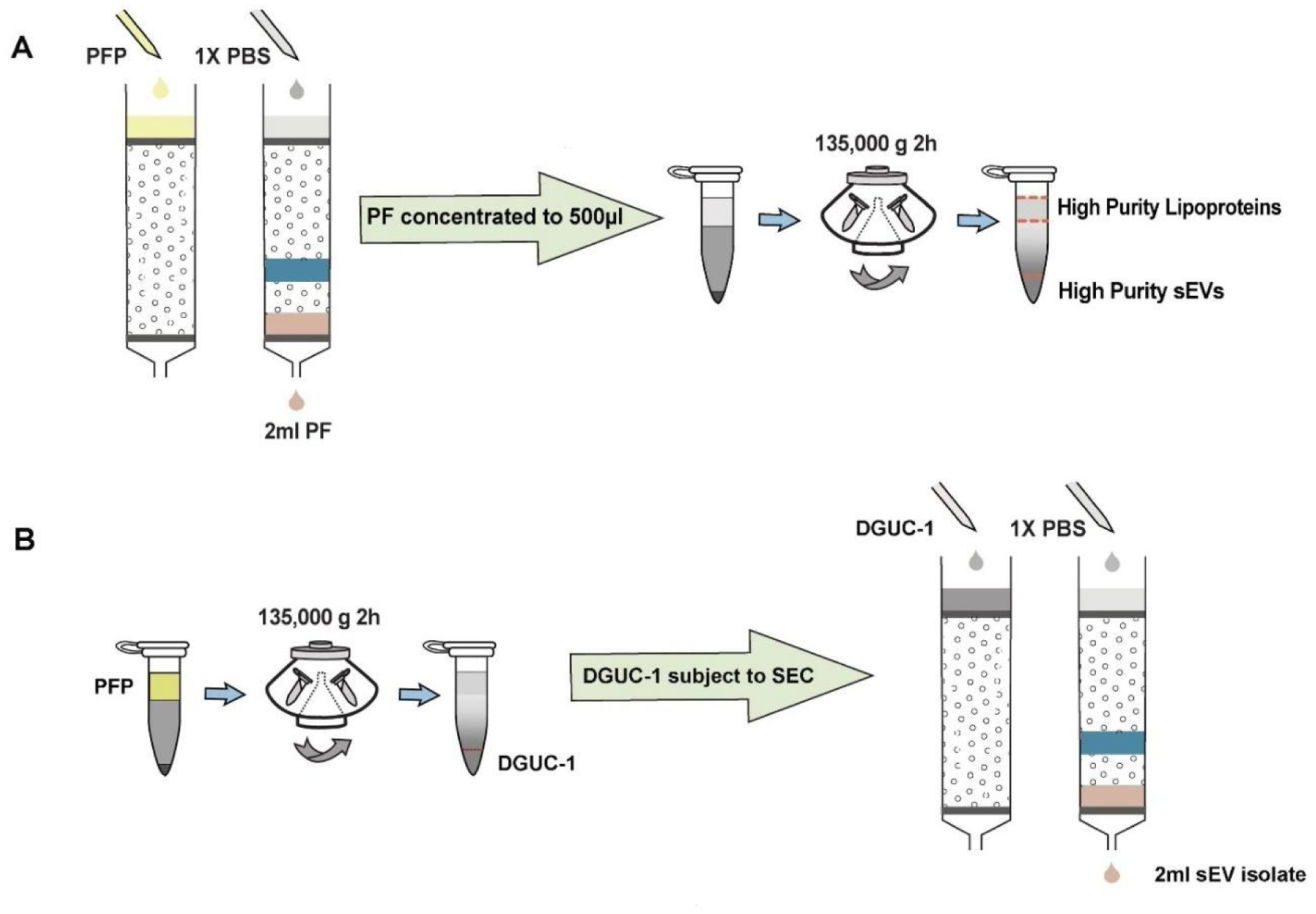
Schematic representation of the SEC-DGUC and DGUC-SEC protocols. (A) Illustrates the SEC-DGUC protocol sequence. Starting from the left, SEC is used initially to separate plasma proteins and HDL. Subsequently, DGUC is employed to separate low-density lipoproteins (i.e., IDL, VLDL, LDL) from sEVs. (B) Depicts the DGUC-SEC protocol, wherein DGUC is initially employed to segregate high-density components from those of lower density, before SEC is applied to partition any remaining proteins from the sEVs.

**Figure 5:**
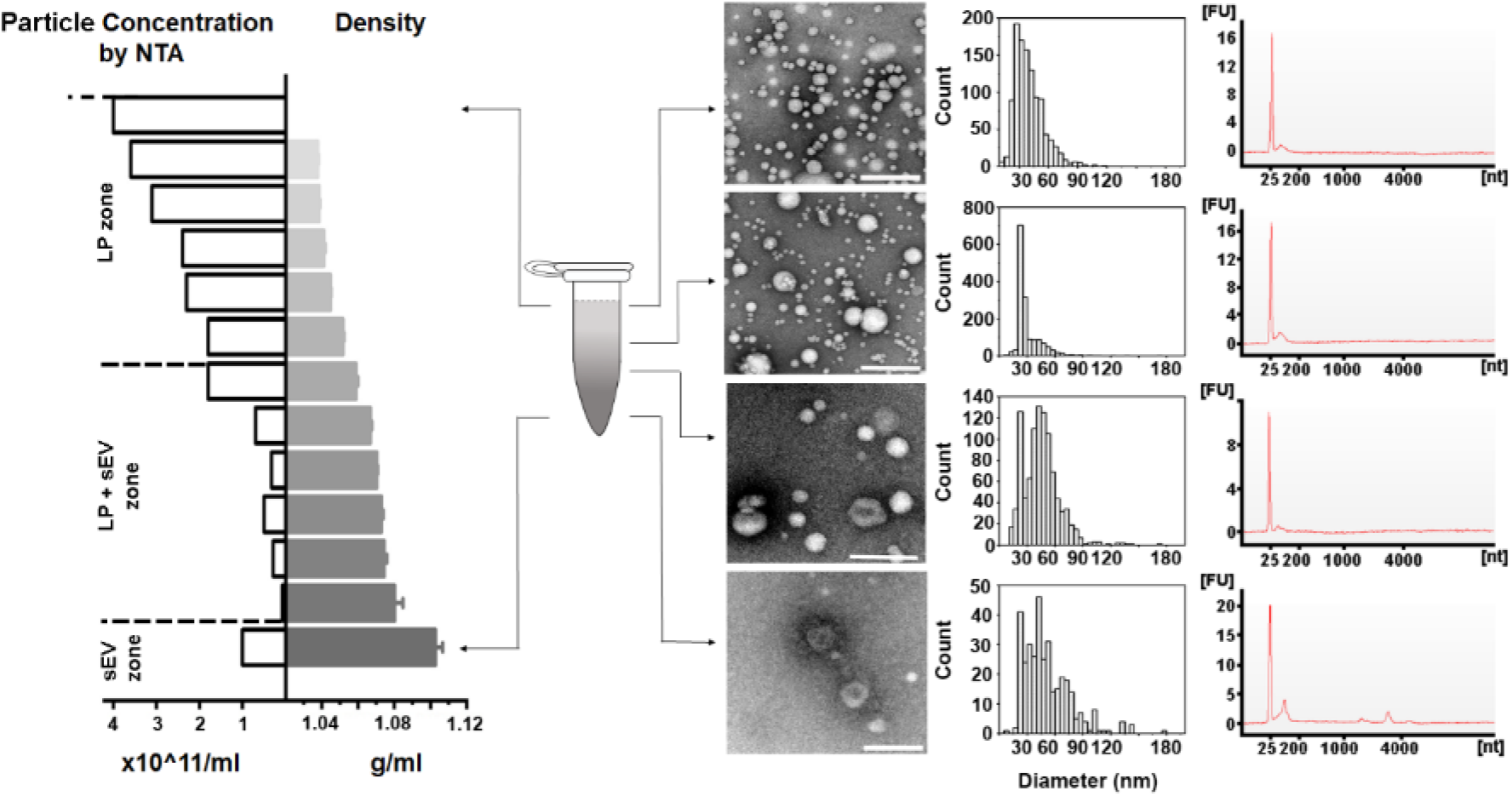
DGUC in the 1.5 *ml* tube format effectively separated sEVs from lipoproteins. PFs obtained from SEC were concentrated into 500 *μl*, loaded onto the density gradient and subjected to ultracentrifugation as previously described. After DGUC, the 1.5 *ml* tube was fractionated into 13 fractions, which were each examined for their particle concentration (by NTA) and the presence of sEVs and lipoproteins (by TEM). In the sEV zone (bottom of the tube), where density was higher than 1.08 *g/ml*, high purity sEVs were indeed observed. A mixed population of sEVs and lipoproteins was observed within the density zone of 1.05∼1.08 *g/ml*. Interestingly, the particle numbers in this density zone were low, thus creating an effective separation zone between lipoproteins and sEVs. The RNA profiles of corresponding particle populations are shown on the right. All scale bars represent 200 *nm*.

To further substantiate the purity of the sEV isolates obtained through SEC-DGUC protocol, we embarked on measuring the total RNA content of the fractions and conducting western blot analysis (see Materials and methods). For these experiments, we simplified the fractionation of the 1.5 *ml* tube into four distinct fractions, denoted as SEC-DGUC-1 to 4 (Figure 5 right side, see Materials and methods). Total RNA analysis revealed that SEC-DGUC-1 had a higher concentration of RNA compared to the other three fractions (Figure 5). The western blot analysis showed an abundant presence of sEV markers such as CD63, CD81, CD9, TSG101, and Flotillin-1 in SEC-DGUC-1, while the lipoprotein marker, ApoB, was notably low. This contrasted with SEC-DGUC-2 to 4, where ApoB was abundant, reinforcing the high purity of sEVs in SEC-DGUC-1. Notably, SEC-PF exhibited a high level of ApoB and low expression of sEV markers (Figure 6B).

**Figure 6:**
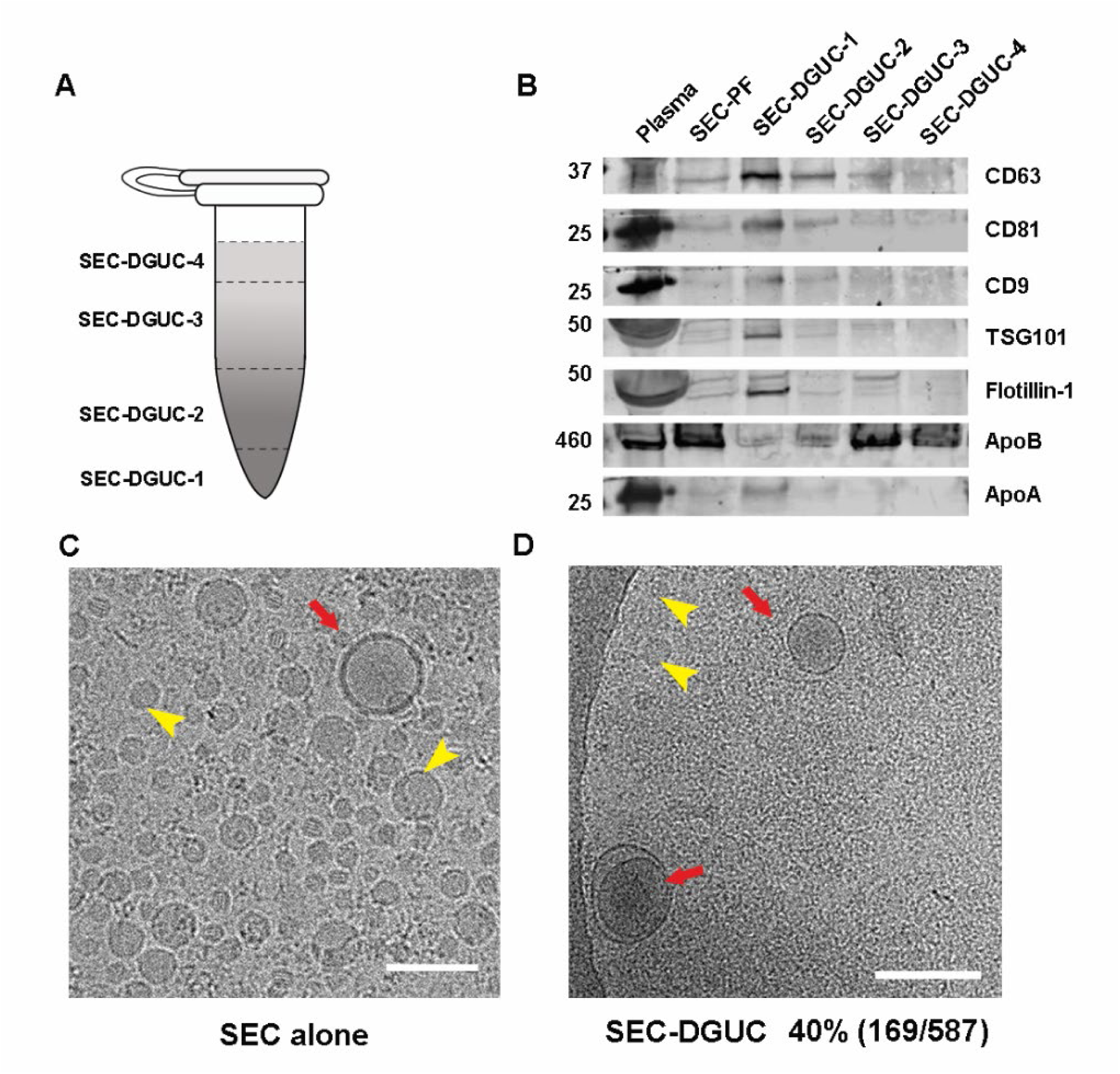
The high purity of sEVs in SEC-DGUC-1 was demonstrated by WB and Cryo-EM. (A) The 1.5 *ml* tube subjected to the SEC-DGUC protocol was fractionated into 4 fractions for easier analysis. (B) Western blot analyses of original plasma, SEC-PF, SEC-DGUC-1 to 4 using sEV and lipoprotein markers. (C) & (D) Cryo-EM images showing the comparison between SEC-PF and SEC-DGUC-1. The SEC-DGUC-1 in (D) was obtained from non-fasting plasma collected in EDTA tubes. All scale bars represent 200 *nm*.

To quantify the purity of sEVs in SEC-DGUC-1, cryo-EM was employed to discern sEVs and lipoproteins (Figure 6D, Supplementary Figure 5). The purity of sEVs, defined as the percentage of sEVs among all particles, was calculated to be 40%, where other 60% particles were mainly small lipoproteins. Considering that SEC-PFs were comprised of only ∼1% of sEVs, the DGUC managed to eliminate over 98% of the lipoproteins.

Interestingly, when the ultracentrifugation time was increased to 16 hours, the size histogram and particle concentration profile closely mirrored those of the 2-hour DGUC (Supplementary Figure 4C-E). This observation suggests that a 2-hour DGUC was sufficient to separate sEVs from lipoproteins in the 1.5 *ml* tube using the fixed-angle rotor.

In conclusion, the SEC-DGUC protocol successfully isolated sEVs from a small volume of plasma under three hours in total by leveraging the strengths of SEC and DGUC.

### 2.5 DGUC-SEC protocol also isolated high-purity sEVs

An alternative approach to the sequential SEC-DGUC process involves reversing the order, implementing DGUC prior to SEC (termed DGUC-SEC) [14,18]. DGUC separates high-density particles from low-density ones in the plasma, while SEC further differentiates proteins and HDL from sEVs. Theoretically, the DGUC-SEC protocol should also effectively isolate sEVs from plasma [19]. However, it remains intriguing to assess if the sequence of applying SEC and DGUC impacts the quality of sEV isolates from a small volume of plasma.

For this purpose, 500 *µl* of plasma was initially processed through a 2-hour DGUC protocol using the 1.5 *ml* tube format. The bottom 120 *µl* high-density fraction (plasma-DGUC-1) was then collected and subjected to SEC (Materials and methods). Similar to the SEC-DGUC protocol, this DGUC-SEC protocol also yielded a high-purity sEV isolate, termed DGUC-SEC-PF, as confirmed by Western blot and transmission electron microscopy (TEM) (Figure 7A, D).

**Figure 7:**
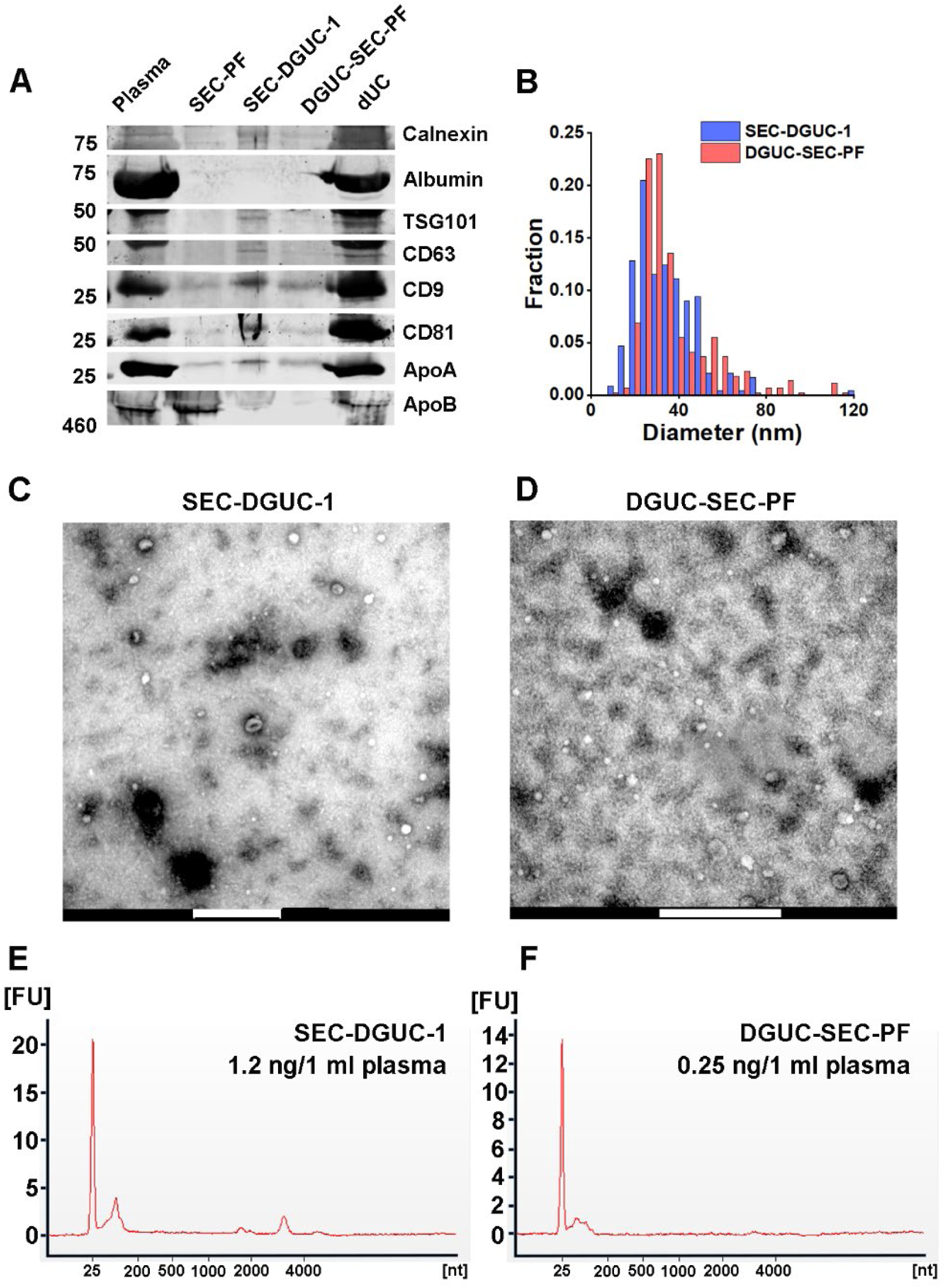
Comparison of sEVs isolated by SEC-DGUC and DGUC-SEC protocols. (A) Western blot of SEC-DGUC-1, DGUC-SEC-PF, sEV isolated by dUC obtained from 2 *ml* of plasma of the same source. (B) Particle size distributions in SEC-DGUC-1 and DGUC-SEC-PF measured by TEM. (C) and (D) TEM images of SEC-DGUC-1 and DGUC-SEC-PF. (E) and (F) Total RNA analyses of SEC-DGUC-1 and DGUC-SEC-PF. Note that data in (E) is the same data shown in the bottom right subfigure of Figure 5.

### 2.6 Comparison of SEC-DGUC and DGUC-SEC protocols: SEC-DGUC obtained higher yield of sEVs

A side-by-side comparison of sEV isolates from the same plasma source using SEC-DGUC and DGUC-SEC was carried out using several independent characterization methods: western blot, TEM and total RNA analysis (Figure 7, see Materials and methods). In the western blot, an additional sEV sample isolated by a routine dUC protocol was included for comparison. Both sEV samples isolated by the combination of SEC and DGUC using the same amount of plasma showed low contaminants (Calnexin, Albumin, ApoA, and ApoB) and clear signal of sEV markers such as CD9, CD63, CD81, and TSG101 (Figure 7A). In stark contrast, sEV isolated by dUC showed a high degree of contamination of Albumin, ApoA, and ApoB, but also an abundance of CD9, CD81, and TSG101. TEM images demonstrated similar appearances of the particles isolated by the two protocols. The particle size distributions of isolated sEVs obtained from the TEM images by both SEC-DGUC and DGUC-SEC protocols also largely overlapped with each other (Figure 7B), further suggesting that the particles isolated by both protocols are similar both in chemical but also in physical aspect. Despite the similarity of SEC-DGUC-1 and DGUC-SEC-PF samples, the SEC-DGUC-1 did show higher signal intensity for all four tested sEV markers (CD9, CD63, CD81, and TSG101) with estimated concentrations approximately 2.1, 2.1, 4.7, and 4.2 times higher compared to the DGUC-SEC-PF in the western blot. In addition, the total RNA analysis (Figure 7E, F) showed that SEC-DGUC-1 contained more than 4 times the total amount of RNA than DGUC-SEC-PF, suggesting that the SEC-DGUC protocol yielded significantly more sEVs.

In conclusion, both the SEC-DGUC and DGUC-SEC protocols proved effective in isolating high-purity sEVs from plasma. Yet, in the context of our experiment, which aimed to isolate sEVs from low plasma volumes, the SEC-DGUC protocol outperformed DGUC-SEC, yielding a significantly greater quantity of sEVs.

### 2.7 High purity of sEVs facilitate downstream analyses

Building upon the results of the comparative analyses, we further examined the sEVs isolated by SEC-DGUC through additional downstream analyses—surface marker detection by flow cytometry (FCM) and protein content analysis by liquid chromatography with tandem mass spectrometry (LC-MS/MS).

The sEV surface proteins were analyzed by FCM using a commercial kit such as the MACSPlex Exosome kit, in a semi-quantitative manner (Figure 8A and B, see Materials and methods). A panel of 37 surface markers comprehensively evaluated the sEVs in the plasma concerning their origin and relative amount.

**Figure 8:**
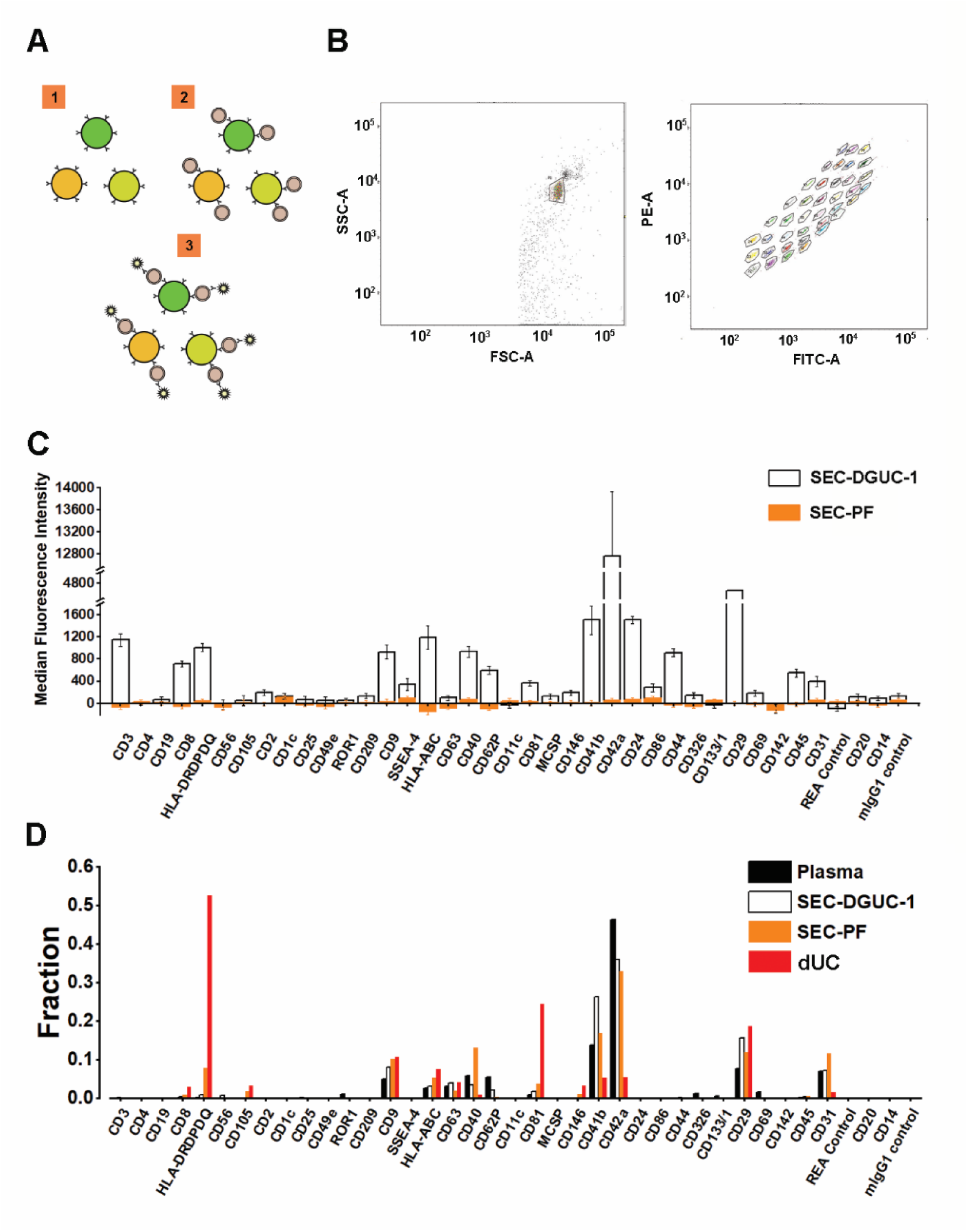
Flow cytometry assay using MACSPlex exosome kit to evaluate sEV isolates. (A) Illustration of the principle of MACSPlex exosome kit. 37 types of beads with different fluorescent colors are functionalized with specific antibodies against sEV surface proteins. sEVs captured by the beads are detected with flow cytometry using APC labeled anti-CD9, CD63, CD81 antibodies. (B) Flow cytometry gating setup. (C) A representative data showing the comparison of median fluorescence intensity (MFI) ± standard error of the mean (SEM) from sEV isolates obtained from SEC-PF and SEC-DGUC-1. Equal numbers of particles (5×10^8^, based on NTA) were loaded for both samples. (D) Comparison of signal patterns among plasma, SEC-DGUC-1, SEC-PF and dUC. The Results showed that sEVs isolated from dUC deviate from plasma whereas, sEVs in the SEC-PF and SEC-DGUC-1 mirror the sEV population in plasma.

We compared the surface marker signals of SEC-DGUC-1 with SEC-PF (Figure 8C), derived from the same plasma source, using the same amount of particles (5×10^8^, calculated from the NTA data), which was reported to be a saturating number [20]. The surface marker panel demonstrated striking differences in signal strength between SEC-DGUC-1 and SEC-PF. For some highly expressed surface markers, such as CD42a, CD41b, and CD29, the signals of SEC-DGUC-1 were up to three orders of magnitude higher than SEC-PF (Figure 8C). Moreover, for other surface markers such as CD3, CD19, HLAs, CD44, CD45, CD31, and SSEA-4, SEC-DGUC-1 provided much higher signals than the background, compared to SEC-PF, which showed undetectable levels of signals. sEV isolates from two more different plasma exhibited a similar trend (Supplementary Figure 6). The contrast between signals from SEC-DGUC-1 and SEC-PF further confirmed the high purity of sEVs in SEC-DGUC-1 and suggested that high-purity sEV isolates would lead to higher sensitivity in the FCM analysis.

Another crucial question to address is whether the sEVs isolated by SEC-DGUC represent the sEV population in the original plasma. To investigate this, we compared the signal patterns of the 37 surface markers across plasma, SEC-PF, SEC-DGUC-1, and sEVs isolated by dUC (Figure 8D). The signal pattern was calculated by normalizing individual surface marker signals to their corresponding signal sum. The number of particles loaded for plasma, SEC-PF, SEC-DGUC-1, and sEV isolated by dUC were 1×10^10^, 1×10^10^, 5×10^8^, and 5×10^8^ (based on NTA measurement), respectively. The number of particles loaded for plasma and SEC-PF was 20 times more than SEC-DGUC-1 to achieve a comparable level of sEV loading. The surface marker signal patterns of SEC-PF and SEC-DGUC-1 were similar to plasma, whereas sEVs from dUC deviated from the plasma (Figure 8D). This suggested that the sEVs isolated by the SEC-DGUC protocol represented the sEV population in plasma, whereas sEVs isolated by dUC represented a biased population.

In summary, SEC-DGUC-1 significantly improved the sensitivity of surface marker detection compared to SEC-PF in flow cytometry, and the sEVs in SEC-DGUC-1 were found to be representative of the sEV population in the original plasma.

Next, the SEC-PF and SEC-DGUC-1 obtained from 5 *ml* of a single plasma source were analyzed by LC-MS/MS (Material and Methods). 343 proteins were identified in SEC-PF, and 754 proteins were identified in SEC-DGUC-1. Out of the 754 proteins identified in SEC-DGUC-1, 51 proteins matched with the top 100 sEV proteins from Vesiclepedia. Specific sEV-associated proteins such as CD81, CD9, and TSG101 were identified. Additionally, other common EV-associated proteins such as 14-3-3 protein (theta, zeta/delta), Rabs (Rab-7A and Rab-8A), ADAM10, EHD4, PFN1, immune system-related proteins (MHC Class II), MSN, SDCBP, ENO1, Annexin, GAPDH, ACTN4, TUBA1C, and ESCRT accessory (Clathrin) were also identified. The following cell-specific markers were detected: CD41, CD36, CD42b for platelets; CD37 for leukocytes, and CD163 for macrophages. Regarding the co-isolated contaminating lipoproteins in the sEV isolate, only ApoA was detected in the top 50 most abundant proteins according to emPAI (Supplementary Table I), which was expected based on the western blot result (Figure 6B). On the other hand, for SEC-PF, 7 out of the top 11 most abundant proteins were lipoproteins (Supplementary Table II). The identified proteins were further analysed with the functional annotation tool on the platform of the Database for Annotation, Visualization, and Integrated Discovery (DAVID v6.842), of which the top 10 in the functional annotation chart were shown in Figure 9. Extracellular exosome was the top first group for SEC-DGUC-1 (Figure 9A), suggesting the identified proteins were highly associated with sEVs. In comparison, the top 3 annotation functions of proteins identified in SEC-PF were not related to sEVs (Figure 9B).

**Figure 9:**
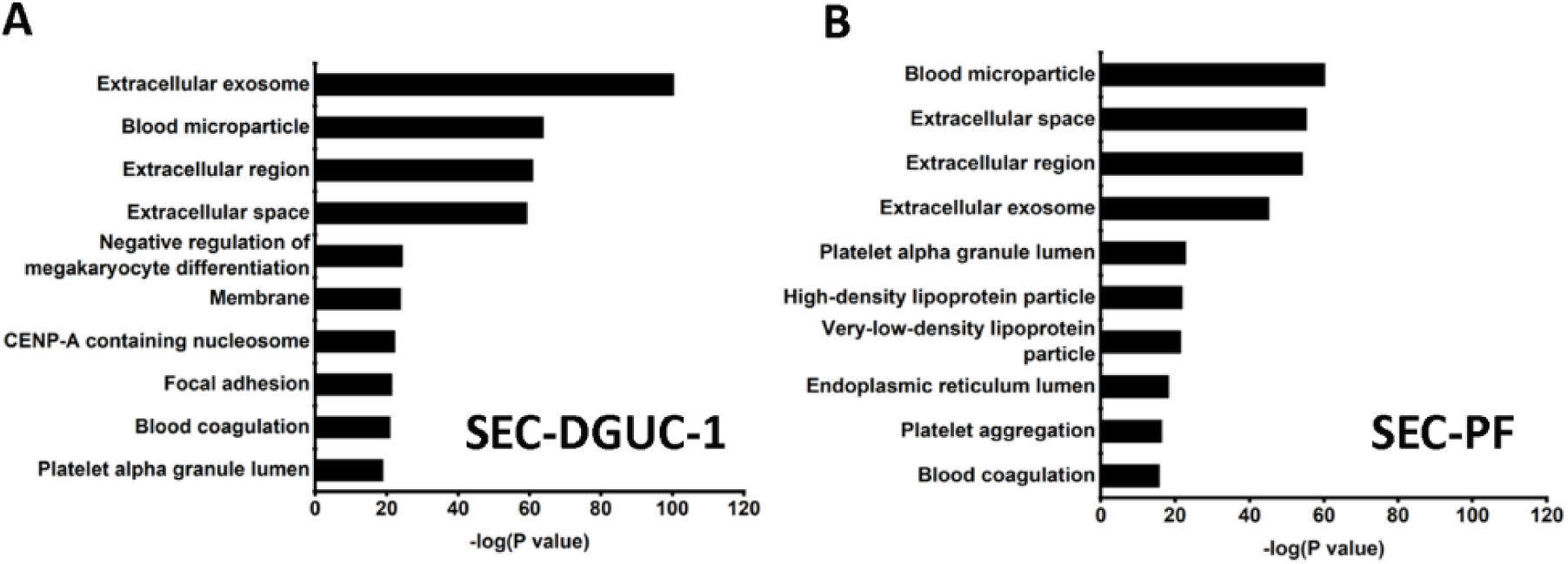
Functional annotation of proteins identified by LC-MS/MS. (A) Functional annotation of proteins identified in SEC-DGUC-1. (B) Functional annotation of proteins identified in SEC-PF.

In summary, sEVs isolated from a small volume of plasma using the SEC-DGUC protocol can provide the required purity to identify sEV-associated proteins in mass spectrometry.

### 2.8 Repeatability and reliability of the SEC-DGUC protocol

We subsequently assessed the repeatability and reliability of the SEC-DGUC protocol in sEV isolation from plasma, critically analysing variations at each step of the protocol.

The repeatability of the SEC step was examined using multiple SEC-PFs derived from the same plasma source. Comparisons were based on particle concentration and size distribution (via NTA) once concentrated to 500 *µl*. This test, utilizing six distinct plasma sources, revealed that the coefficient of variation (CV) in particle concentration ranged between 6% and 55% (Supplementary Figure 7 table). Meanwhile, size distributions exhibited no notable variations across plasma samples (Supplementary Figure 7A-C), implying that despite potential variations in particle numbers, the particle populations remained largely consistent across PFs. The complete SEC-DGUC protocol was then scrutinized for its repeatability using multiple sEV isolates (SEC-DGUC-1) from the same plasma source. Particle concentrations and size distributions were evaluated, yielding CVs of 24% and 25% for particle concentration in SEC-DGUC-1 (Supplementary Figure 8 table). We then compared size distributions for each plasma using Jensen-Shannon Divergence (JSD) in each fraction. The JSD values are well below 0.1 (Figure 10B), indicating a consistent population of isolated particles, as further supported by Supplementary Figure 8. The repeatability of the SEC-DGUC protocol was further evidenced by its consistent particle concentration profile across the 1.5 *ml* tube. Four technical replicates of the profiles showed good repeatability (Figure 10A), suggesting the SEC-DGUC protocol’s ability to robustly generate a consistent density gradient profile and particle subpopulations. Additionally, NTA-measured size distributions displayed well-overlapped histograms of particles (Figure 10B), reinforcing the protocol’s robustness.

**Figure 10:**
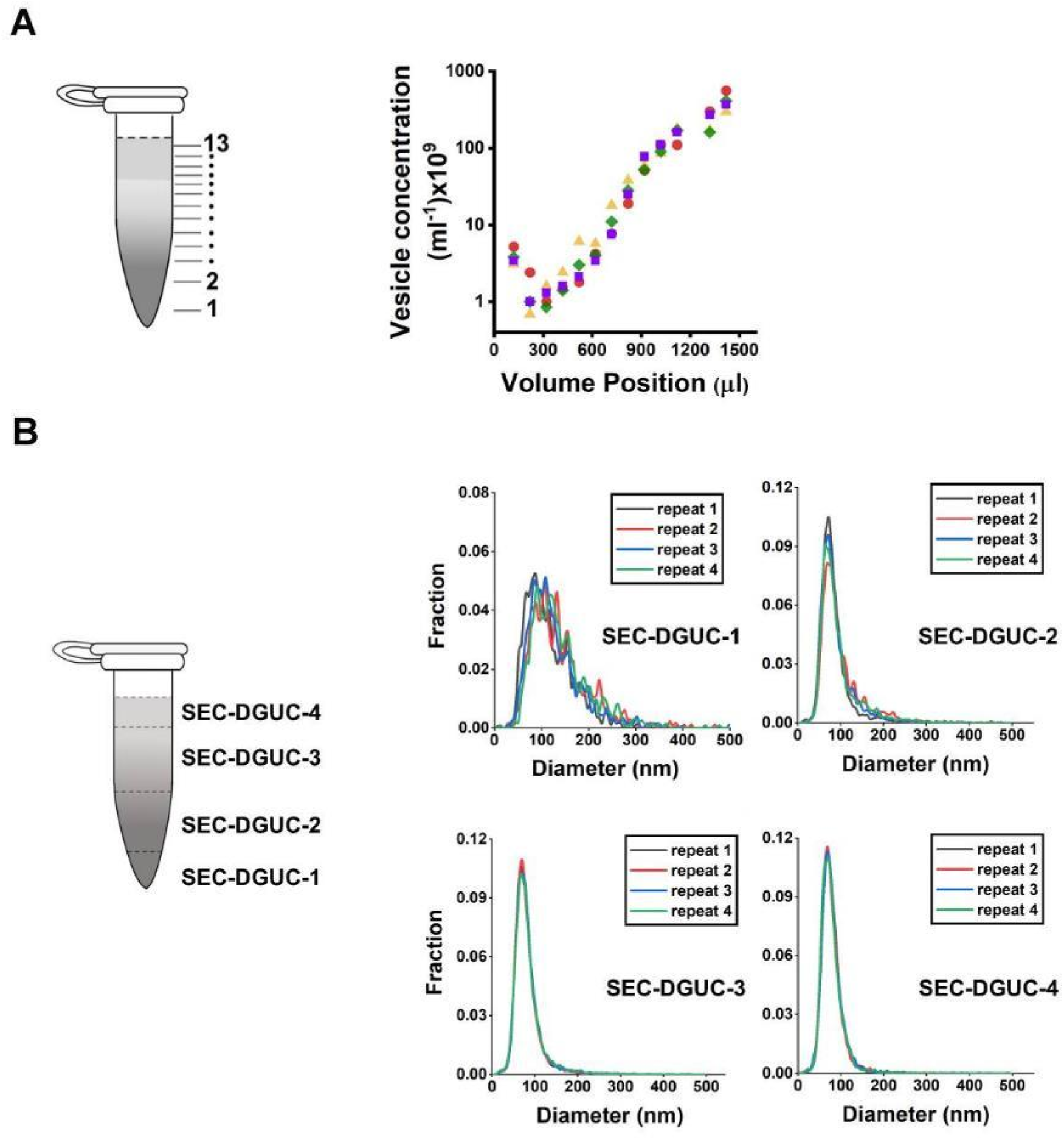
Repeatability of the SEC-DGUC protocol. (A) Particle concentration profiles (by NTA) along the 1.5 *ml* tube from 4 technical replicates subjected to SEC-DGUC protocol. (B) Particle size distributions measured by NTA in the four fractions indicated on the left figure. Four technical replicates subjected to SEC-DGUC protocol were shown. JSD values for SEC-DGUC-1 to 4 are 0.015, 0.006, 0.001, and 0.002, indicating strong similarities among the histograms.

In the subsequent phase, we gauged the protocol’s reliability across various plasma samples by evaluating particle concentration profiles, size distributions, and TEM images of SEC-DGUC-1. Particle concentration profiles derived from diverse plasma samples displayed a signature ‘tick’ shape, indicative of reliable sEV and lipoprotein separation irrespective of the plasma source (Supplementary Figure 9). However, the size distributions of SEC-DGUC-1 varied across different plasma samples (Supplementary Figure 10), suggesting that the characteristics of sEVs were plasma-specific. TEM images of SEC-DGUC-1 derived from five different biobanked plasma samples further confirmed the high purity of sEVs, with many particles exhibiting the characteristic cup-shape and a low contrast (Figure 11).

**Figure 11:**
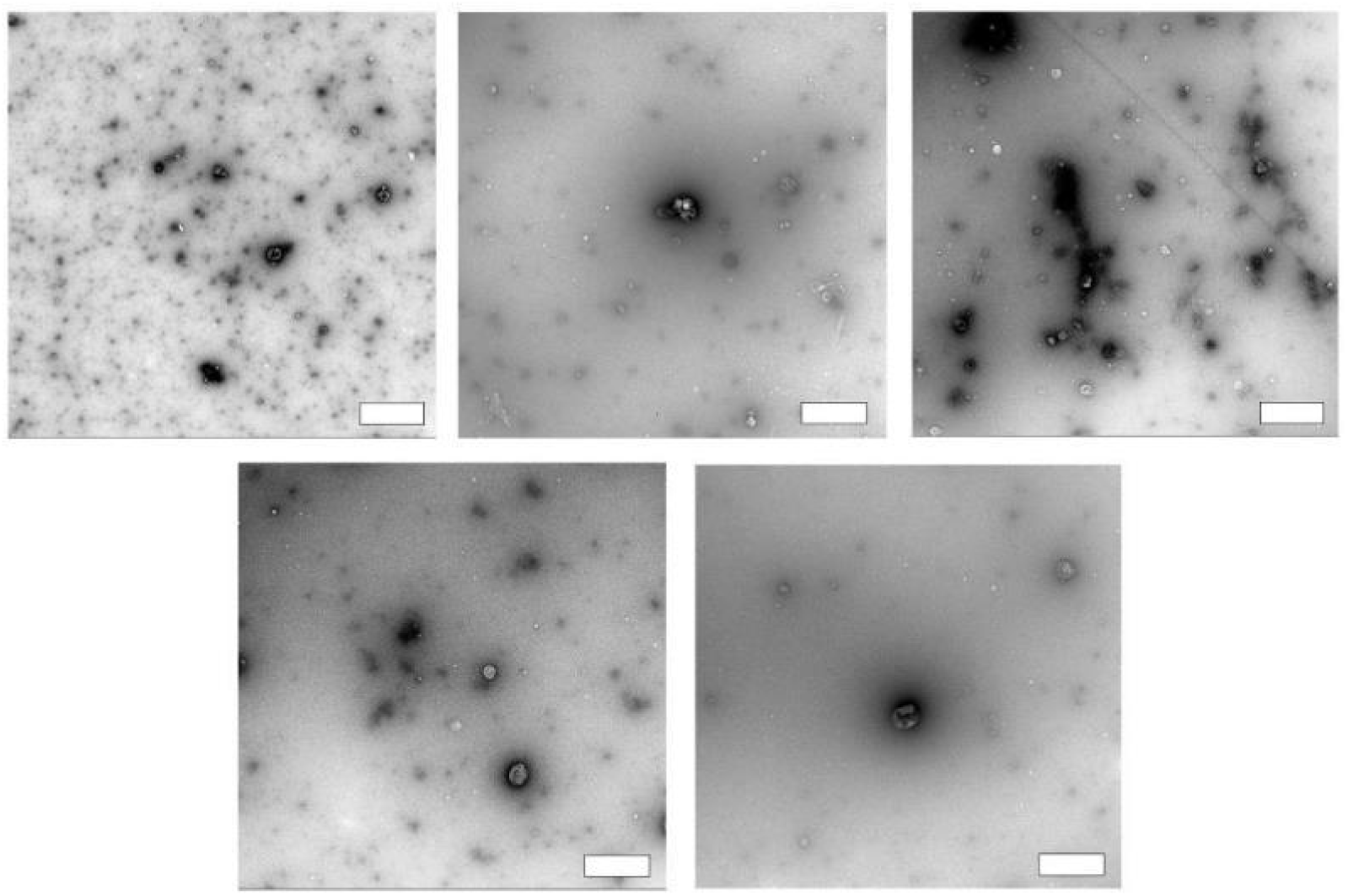
Reliability of the SEC-DGUC protocol. TEM images of SEC-DGUC-1 were obtained from 5 fasting plasma (Biobank samples, EDTA tubes). High purity of sEVs represented by the low-contrast, cup-shaped vesicles was evident across different samples, suggesting the SEC-DGUC was reliable in obtaining high-purity sEVs. All scale bars represent 200 *nm*.

**Table I.** Particle numbers in different steps during the SEC-DGUC protocol, calculated based on NTA measurement. The first column provides the total particle numbers measured in 500 *μl* fasting plasma (∼1 *ml* of whole blood) from 5 individuals. The second column provides the corresponding total vesicle numbers in SEC-PF after the plasma was subjected to SEC. The third column provides the corresponding total vesicle numbers in SEC-DGUC-1 after the SEC-PF was subjected to DGUC.

| Plasma | SEC-PF | SEC-DGUC-1 |
| --- | --- | --- |
| $8 \times 10^{10}$ | $6.5 \times 10^{10}$ | $1.8 \times 10^9$ |
| $5.5 \times 10^{11}$ | $1.65 \times 10^{11}$ | $1.32 \times 10^9$ |
| $1.75 \times 10^{11}$ | $8.5 \times 10^{10}$ | $1.68 \times 10^9$ |
| $6 \times 10^{11}$ | $1.2 \times 10^{11}$ | $1.56 \times 10^9$ |
| $2.4 \times 10^{10}$ | $1.6 \times 10^{10}$ | $1.2 \times 10^9$ |

In conclusion, the SEC-DGUC protocol demonstrated a CV of ∼25% for particle number repeatability when isolating sEVs from the same plasma source. Despite this, it exhibited a reliable capacity to isolate high-purity sEVs from varying plasma sources.

## 3. Discussion

The task of isolating small extracellular vesicles (sEVs) from human plasma becomes particularly challenging when dealing with minimal volumes. To tackle this challenge, our study scrutinized two commonly employed methods –SEC and DGUC, and further investigated the efficiency of their combined use. We found that neither SEC nor DGUC, when used individually, provide an optimal solution for sEV isolation. SEC, although effective in removing plasma proteins, results in a particle fraction dominated by lipoproteins (99%). DGUC, on the other hand, facilitates substantial particle and protein removal due to its density-based separation; however, it does not completely eliminate protein contamination.

The joint use of these methods has been explored previously, yet for small plasma volumes, understanding how to strategically integrate these techniques to maximize efficiency and sEV purity is crucial. Our devised SEC-DGUC protocol merges the advantages of both techniques, employing SEC to eliminate plasma proteins and high-density lipoproteins (HDL), and DGUC to remove remaining lipoproteins.

Interestingly, our results showed that performing SEC prior to DGUC yields a higher yield of sEVs, especially for plasma volumes ranging from 500 *µl* to 2 *ml*. The gel matrix of the SEC column tends to retain a portion of proteins and particles, causing sample loss. When a low particle concentration sample is processed through an SEC column, this loss becomes more significant. Consequently, performing SEC first—when sEVs, lipoproteins, and plasma proteins are all present—distributes the loss among all components. Given the minor proportion of sEVs in plasma, the loss is more likely absorbed by lipoproteins and plasma proteins. Conversely, protocols that use SEC in the final isolation step could experience a significant loss of sEVs, as most lipoproteins and plasma proteins have already been removed [12]. Such protocols may yield a lower amount of sEVs or require a larger plasma volume to obtain enough sEVs for subsequent analyses [12,14]. This could explain the lower yield of sEVs observed in the DGUC-SEC protocol compared to the SEC-DGUC protocol.

The SEC-DGUC strategy has been adopted by others and has shown reliable results [13]. However, DGUC is typically performed in tubes of larger volume capacity, such as 12 *ml* [13] or 16.8 *ml* [14] placed in swinging-bucket rotors. For our SEC-DGUC protocol, the DGUC was carried out in a 1.5 *ml* tube format and the density gradient setup was designed to harvest all sEVs with a density greater than 1.08 *g/ml*, rather than in a specified narrow range of density. From our experiments, we recognized that small tubes are easier to handle, require shorter handling time as well as lower volumes of reagents, are more suitable for benchtop ultracentrifuges, and the fractions can be harvested manually without the need of specialized equipment. More importantly, DGUC with the small-tube format is done using a fixed-angle rotor which has a shorter sedimentation path length. This requires a much shorter time to isolate sEVs (< 3 hours) compared to a larger volume tube format using a swinging-bucket rotor which typically lasts ∼ 16 hours or more [21]. Moreover, DGUC using the small-tube format resulted in high-purity sEV isolates collected in a volume of 120 *µl*, the concentration of which enabled direct downstream analyses such as electron microscopy, western blot, total RNA analysis and a multiplex bead-based flow cytometric assay.

The advantage of high-purity sEV isolates was demonstrated by flow cytometry and mass spectrometry. Flow cytometry result demonstrated that the sEVs isolated by SEC-DGUC gave a signal 3 orders of magnitude higher than using SEC alone, thereby improving the sensitivity of the assay to a greater extent. Moreover, the surface marker pattern of 37 proteins helps to assess the nature of the sEV isolates. The surface marker patterns matched closely among the sEVs isolated from SEC-DGUC, SEC alone and the original plasma, suggesting a close representation of sEVs isolated from SEC-DGUC to the original plasma. It is important to note that, the surface marker patterns of sEVs isolated from a routine dUC protocol deviated significantly from sEV populations in the original plasma.

There have been several studies using mass spectrometry to determine the proteome of plasma-derived EVs [14,22–24]. Karimi et al. 2018 analysed sEVs isolated by a DGUC-SEC protocol from large volumes of pooled healthy human plasma with LC-MS/MS and identified 1187 proteins. Many proteins identified were sEV-associated proteins such as CD63, CD9 and CD81 and the identified proteins were closely associated with extracellular exosomes. In comparison, when analysing the sEVs isolated by the newly introduced SEC-DGUC protocol using 5 *ml* plasma from a single healthy individual, the proteomics data identified 753 proteins which included sEV-specific proteins, such as CD81, CD9, Flotillin and TSG101. Moreover, the proteins were closely associated with extracellular exosomes as well. We expect that, with a better optimized LC-MS/MS protocol, the volume of plasma required for the sEV isolation can be further reduced. Therefore, we conclude that the sEVs isolate obtained from the small volume plasma SEC and DGUC protocol is of sufficient yield and purity for mass spectrometry analysis.

The key to the consistency of the SEC-DGUC protocol lies in three aspects: first, the capability of SEC to efficiently remove plasma proteins and HDL; second, the capacity of DGUC to separate sEVs from low density lipoproteins; third, the careful manual handling during DGUC, especially in the final step of harvesting the sEVs from the tube. When evaluated individually, SEC is a well-established and quick method which is commercially available; DGUC with the simple density gradient design enables consistency, precision and the potential to fully automate the protocol thereby mitigating human error. For these reasons, the SEC-DGUC protocol can be extrapolated to obtain high-purity sEVs in a repeatable and reliable manner. Collectively, our results clearly demonstrated the quality of the isolated sEVs and the robustness of SEC-DGUC protocol in a small-tube format. This is not to claim that the existing protocols are less reliable than the currently developed protocol. In fact, there is room to further improve the protocol in future efforts.

The SEC-DGUC protocol, though efficient, has limitations like potential damage to or aggregation of sEVs [25], and low-level contamination with HDL. The iodixanol reagent introduced in DGUC could potentially interfere with certain downstream analyses, and variations in plasma samples might impact sEV purity. Although the protocol effectively isolates representative sEVs from plasma, some lower density sEVs that require a longer centrifugation time may be missed.

Isolating high-purity sEVs from small volumes of plasma is essential in fully realizing the potential of using sEVs as biomarkers. It is of particular importance to standardize the sEV isolation protocol so that the data obtained by different labs can be compared and integrated with high confidence. The SEC-DGUC protocol presented here is relatively short, easy to carry out, and isolates sEVs with high purity in a repeatable and reliable manner. The SEC-DGUC protocol can be quickly adapted by any laboratory and can aid clinical studies that have limited accessible plasma volumes. It potentially offers a pathway for the standardization of sEV isolation from plasma.

## 4. Materials and methods

### Blood collection and preparation of platelet-free plasma

All procedures involving peripheral blood specimens were approved by the Singapore National Health Group Domain Specific Review Board (the central ethics committee) and were mutually recognized by the Nanyang Technological University Institutional Review Board (IRB#2018/00671). All blood specimens were de-identified prior to their use in the experiments. The plasma used in this study was obtained from two sources, fresh blood and biobanked plasma. Fresh blood was obtained with written informed consent from healthy adult males aged between 25∼45 years. All blood was collected after overnight fasting unless specified otherwise in the figure legend. The volunteers were free of medications for at least 3 weeks. Blood was drawn in the morning using venepuncture technique with a 21G butterfly needle into anticoagulant citrate dextrose-A (ACD-A) containing vacuette tubes (455055, Greiner, Austria). Three vacuette tubes (each 9 *ml* volume) of blood was collected from each volunteer. The vacuette tubes were inverted a few times to mix the blood with the anticoagulant, placed in an upright position and processed within 20 minutes of collection. We applied the International Society on Thrombosis and Haemostasis (ISTH) protocol to prepare the platelet-free plasma (PFP). Briefly, the blood was centrifuged at 2500 *g* for 15 min at room temperature using a table-top centrifuge (Z206A, Hermle Labortechnik GmbH, Germany) to remove blood cells, resulting in platelet-poor plasma (PPP). The PPP was transferred into a new tube and centrifuged once again at 2,500 *g* for 15 minutes, resulting in PFP. The PFP was transferred into a new tube, homogenized by gentle inversion, split into 500 *μL* aliquots in 1.5 *ml* eppendorf tubes and stored at -80 °C until further use. The biobanked plasma samples were obtained from healthy males following overnight fasting with written informed consent. Briefly, the blood was collected using venepuncture technique with a 21G butterfly needle into ethylenediaminetetraacetic acid (EDTA) containing vacuette tubes. The blood was then centrifuged at 1200 *g* for 15 *min* at 4°C with brake-two in a horizontal swing-bucket centrifuge. The plasma fraction was collected within 0.1 *ml* of the interphase with the buffy coat layer. The plasma was split in 500 *μl* aliquots in 1.5 *ml* eppendorf tubes and stored at -80 °C until further use. All data shown in this paper used ACD-A plasma unless indicated otherwise in the figure legends.

### Size exclusion chromatography

An aliquot of frozen plasma was taken from –80 °C and allowed to thaw completely at room temperature. SEC elution was performed using PURE-EVs SEC columns (HBM-PEV, HansaBioMed Life Sciences, Tallinn, Estonia) following the manufacturer’s protocol. Briefly, the SEC columns were equilibrated to room temperature and washed with 3 x 10 *ml* particle-free 1X phosphate buffer saline (PBS) (SH30256.01, Hyclone, UT, USA) to eliminate preservative buffer residues. A volume of 500 *µl* of PFP was applied on top of each column. Once the sample was inside the gel matrix, the column was loaded with particle-free 1 X PBS (the mobile phase of the SEC column). The column was not allowed to dry out during this process. Each of the fifteen fractions of 500 *µl* volume was collected, after which the column was washed with approximately 20 *ml* of 1 X PBS and stored at 4°C. These SEC columns were washed and reused up to 5 times.

The first six SEC fractions (3 *ml*) constituted void volume, hence were discarded and the rest of the nine fractions were analysed for particle numbers and protein concentration. Fractions 7 to 10 which were subsequently identified as the particle fractions (PF) were pooled to a volume of 2 *ml* and concentrated using an Amicon Ultra 100 kDa centrifugal filter (Merck Millipore, MA, USA), made from regenerated cellulose, to a final volume of 500 *µl*.

### Density gradient ultracentrifugation

To prepare the density solutions, a 50% Working Solution was firstly made by diluting 5 vol of iodixanol (Optiprep^TM^, D1556, Sigma Aldrich, MO, USA) with 1 vol of 0.25 *M* Sucrose, 6 *mM* EDTA, 60 *mM* Tris-HCl (pH 7.4). The Working Solution was then mixed with Homogenization Medium composed of 0.25 *M* sucrose, 1 *mM* EDTA, and 10 *mM* Tris-HCl, (pH 7.4) to prepare density solutions of desired percentages.

For DGUC experiment using the 12 *ml* tube, PFs obtained from 6 *ml* of plasma (24 *ml* total volume as previously described) were concentrated to 6 *ml* and added to a Beckman Coulter ultra-clear centrifuge tube (344059, Beckman Coulter, USA). A density gradient was carefully constructed by underlying 2 *ml* each of 10%, 30%, and 50% iodixanol cushions sequentially. The gradient tube was balanced and subjected to ultracentrifugation at 150,000 x *g* for 2 *h* at 4 *°C* using a swinging-bucket rotor (SW 41 Ti Beckman Coulter, USA).

For the DGUC using the 1.5 *ml* tube format, 500 *µl* of the concentrated PF obtained from a single SEC (as previously described) was added to a Beckman Coulter ultracentrifuge tube (357448, Beckman Coulter, USA). For plasma volume higher than 500 *µl*, SEC-PFs from multiple SECs were pooled and concentrated to 500 *µl* before adding to the ultracentrifuge tube. A density gradient was carefully constructed by underlying 800 *µl* of 10% iodixanol solution and 20 *µl* of 50% iodixanol sequentially. The gradient tubes were balanced and subjected to centrifugation at 135,000 x *g* for 2 *h* at 4 *°C* using a fixed-angle rotor (TLA-55, Beckman Coulter, USA). After DGUC, thirteen fractions starting from the bottom of the tube were manually collected by using gel loading pipetting tips. In total RNA analysis and western blot experiments, fractions 2-4; 5-10 and 11-13 were pooled together for easier analyses. Thus, four main fractions were analysed namely: SEC-DGUC-1 (120 *µl* volume), SEC-DGUC-2 (2-4 pooled, 300 *µl* volume), SEC-DGUC-3 (5-10 pooled, 600 *µl* volume) and SEC-DGUC-4 (11-13 pooled, 300 *µl* volume) (Figure 6A).

### DGUC-SEC

For the DGUC-SEC protocol, 500 *µl* of plasma was added to Beckman Coulter centrifuge tubes (357448, Beckman Coulter, USA) and followed by underlying 800 *µl* of 10% iodixanol solution and 20 *µl* of 50% iodixanol sequentially. The gradient tubes were subjected to centrifugation at 135,000 x *g* for 2 *h* at 4 *°C* using a fixed-angle rotor (TLA-55, Beckman Coulter, USA). After DGUC, the bottom 120 *µl* was collected as the high density fraction and designated as plasma-DGUC-1. The plasma-DGUC-1 was then topped up to a volume of 500 *µl* with 1X PBS to run through the SEC column. Fractions 7 to 10 from the SEC were collected, pooled together, and concentrated to a final volume of 120 *μl* using an Amicon Ultra 100 kDa centrifugal filter (Merck Millipore, MA, USA). For plasma volume higher than 500 *μl,* multiple tubes were processed in parallel in the DGUC step, and the resulting plasma-DGUC-1 fractions were pooled and subjected to SEC to obtain sEV isolates.

### Total protein analysis

Protein contents of the SEC fractions were measured using a BCA protein assay kit (23225, Pierce, Thermo Fisher Scientific, MA, USA). A standard curve (range 0–1000 *μg/ml*) was derived with six points of serial dilution with bovine serum albumin (BSA). BSA standard or samples (10 *µl*) were transferred to a 96-well plate (Thermo Scientific Nunc MicroWell) to which 100 *µl* working reagent was added (working reagent 50:1 ratio of assay reagents A and B). The plate was incubated for 30 min at 37°C before being analysed with a multi-well spectrophotometer at 562 *nm* (Tecan Infinite M200Pro, Switzerland). The average 562 *nm* absorbance measurement of the blank replicates was subtracted from all other individual standard and unknown sample replicates. A standard curve was obtained from the measurement of BSA standard. The protein concentration of each unknown sample was calculated based on the standard curve.

### Density measurement

The density of fractions obtained from DGUC was determined by absorbance spectroscopy (Tecan Infinite M200Pro, Switzerland) reading at 340 *nm* in a flat-bottomed 96-well polystyrene plate (167008 Nunclon, ThermoFisher Scientific, MA, USA). A range of iodixanol solutions (5–50%) was diluted at a 1:1 ratio with deionized water and measured in the same 96-well plate to serve as the standard control [26]. For iodixanol concentrations above 35%, a second dilution of the solutions was done to avoid absorbance values above 1.2. The absorbance measurements were made against water and 0.25 M sucrose blanks. Each sample was prepared by triplicate to reduce error . Prior to measuring the absorbance, the plate was shaken thoroughly to mix the samples. A standard curve was prepared by plotting the predetermined densities [27] of the 2-50% iodixanol solutions against their mean absorbance. Using the standard curve, the densities of the fractions collected from the iodixanol gradient following DGUC were calculated.

### Nanoparticle tracking analysis

Nanoparticle tracking analysis (NTA) was carried out using ZetaView® (PMX 120, Particle Metrix, Meerbusch, Germany). Samples were diluted to the recommended particle concentration with particle-free 1 X PBS prior to analysis. The particle motion was measured over 11 positions with two repeats at each position with the following parameters, frame rate: 30 fps; number of frames recorded: 60; sensitivity: 80; exposure: 100; minimum brightness: 30; minimum pixel size: 10; maximum pixel size: 10000; trace length: 30; temperature: 25 *°C*.

### Western blotting

In the first experiment (Figure 6B), SEC-PF and sEV isolates from 2 *ml* of plasma obtained by SEC and SEC-DGUC respectively were analysed for the presence of sEV and lipoprotein markers. The same loading volumes of 22 *µl* were used for SEC-DGUC-1 to 4. Since 500 *µl* SEC-PF was loaded on top of 820 *µl* iodixanol in the 1.5 *ml* tube, resulting in a total volume of 1320 *µl*, the equivalent loading volume for SEC-PF was calculated to be 22 x 500/1320 = 8.3 *µl*. The loading particle number (based on NTA measurement) of SEC-PF and SEC-DGUC-1 to 4 were 1.8×10^10^, 8.8×10^8^, 5.9×10^8^, 5.5×10^9^, 2.86×10^10^, respectively. In this manner, not only the amount of sEVs against lipoproteins of each fraction can be evaluated, but also the particle concentration profile in the 1.5 *ml* tube after SEC-DGUC can be analysed.

In the second experiment (Figure 7A), SEC-PF and 3 sEV isolates obtained from 2 *ml* plasma using SEC-DGUC, DGUC-SEC and dUC were examined for the presence of sEV and lipoprotein markers. The sEVs from SEC-DGUC, DGUC-SEC and dUC were collected in a volume of 120 *µl* and same loading volumes of 30 *μl* were used. The sample loading volume for SEC-PF (11.4 *µl)* was calculated as described above.

All samples were lysed in radioimmunoprecipitation assay (RIPA) buffer (89900, Pierce, Thermo Scientific, MA, USA) containing protease inhibitor cocktail (36978, Thermo Scientific, MA, USA) while chilling on ice. After mixing with loading buffer (4 x) containing β-mercaptoethanol (80570, Merck, Darmstadt, Germany) and heated to 70 ^°^*C* for 10 *min*, the samples were loaded on 12% SDS-PAGE gels and electrophoresed to detect sEV markers (CD63, CD9, CD81, and TSG101 tested in both Figure 6B and Figure 7A, while Flotillin-1 was tested in Figure 6B but not in Figure 7A), lipoprotein markers (ApoA-I and ApoB), and negative control markers (Calnexin and Albumin, both tested in Figure 7A but not in Figure 6B). After being transferred to 0.45 *µm* nitrocellulose membranes (1620115, Bio-rad, Feldkirchen, Germany), proteins were stained with REVERT total protein stain (926-11015, Li-COR Biosciences, Lincoln, NE, USA) for normalisation. After this, membranes were blocked at room temperature for 1 *h* with Odyssey blocking buffer TBS (927-50000, Li-COR Biosciences, Lincoln, NE, USA), and incubated overnight at 4 *°C* with 1:1000 of the following antibodies: mouse monoclonal anti-CD63 (MX-49. 129.5) (sc-5275, Santa Cruz Biotechnology, Dallas, TX, USA), rabbit monoclonal (9EPR2949) anti-CD9 (ab92726, Abcam, Cambridge, MA, USA); mouse monoclonal anti-CD81 (B-11) (sc-166029 Santa Cruz Biotechnology, Dallas, TX, USA), rabbit polyclonal anti-Flotillin-1 (ab41927, Abcam, Cambridge, MA, USA), rabbit polyclonal anti-TSG101 (ab30871, Abcam, Cambridge, MA, USA), mouse monoclonal anti-ApoA-I (B-10) (sc-376818, Santa Cruz Biotechnology, Dallas, TX, USA), mouse monoclonal anti-ApoB (C1.4) (sc-13538, Santa Cruz Biotechnology, Dallas, TX, USA), mouse monoclonal anti-Albumin (ab10241 Abcam, Cambridge, MA, USA) and rabbit polyclonal anti-Calnexin (ab22595, Abcam, Cambridge, MA, USA). The membranes were washed by TBS with 0.1% Tween 20 (TBS-T), and incubated at room temperature for 1 *h* with IgG secondary antibodies at 1:15,000, i.e., IRDye 800CW anti-mouse (925-32210, Li-COR Biosciences, Lincoln, NE, USA) and IRDye 680LT anti-rabbit (926-68021, Li-COR Biosciences, Lincoln, NE, USA). Membranes were washed again with 1X TBS-T and scanned with an Odyssey CLx imaging system (Li-COR Biosciences) using 700 and 800 *nm* channels. Visualization was done by ImageStudio software version 5.2 (LI-COR Biosciences).

### Transmission electron microscopy

The transmission-electron microscopy experiments were carried out at Cryo-Electron Microscopy Platform, NTU Institute of Structure Biology. A carbon-coated grid (CF300-CU, Electron Microscopy Sciences, Hatfield, PA, USA) was glow-discharged for 1 *min* before placing 4 *µl* sample for adsorption of 1 *min*. After blotting, the grid was negatively stained for 1 *min* using 4 *µl* of 2% uranyl acetate. The grid was then blotted and air-dried for TEM imaging. Grids were imaged using a T12 transmission electron microscope operating at 120 *kV* with an Eagle 4k HS camera. TEM images were analysed using ImageJ [28] with Nanodefine plug-in [29] to identify and count particles. Particles with low contrast were difficult to recognize by the Nanodefine plug-in and they were counted and measured manually using ImageJ measuring tools.

### Cryo electron microscopy

The cryo-electron microscopy experiments were carried out at Cryo-Electron Microscopy Platform, NTU Institute of Structure Biology. SEC-PF and SEC-DGUC-1 samples that demonstrated an NTA concentration of at least 3×10^11^ particles were selected for Cryo-EM imaging. For SEC-DGUC-1, an additional step of dialysis using Pur-A-Lyzer^TM^ Mini 6000 dialysis kit (PURN60030, Sigma-Aldrich, MO, USA) was added to remove iodixanol before grid preparation. The grid preparation protocol follows Wang et al. 2020 [30]. Briefly, a volume of 2∼3 *µl* of the sample was applied to a glow discharged Quantifoil R2/2 grid coated with 2 *nm* carbon (Jena, Germany) and incubated for 5∼10 *mins*. The grid was then loaded into FEI vitrobot. Another 2 *µl* of the sample was added to the grid and immediately blotted with following parameters, blotting time: 3 *s*; humidity: 100%; temperature: 4 *°C*; blotting force: - 1; waiting time: 0 *s*;. and thereafter grids were flash frozen in liquid N2-cooled liquid ethane. Grids were imaged on an Arctica transmission electron microscope (FEI) operated at 200 *kV* on a Falcon III (FEI) direct electron detector. The Cryo-EM images were analysed manually using ImageJ.

### Total RNA isolation

For total RNA isolation, 100 *µl* isolates obtained from 2 *ml* of plasma following the SEC-DGUC (SEC-DGUC 1-4) and DGUC-SEC protocols were used. RNA extraction was done using the miRNeasy Serum/Plasma Kit (217184, Qiagen, GmbH, Hilden, Germany) according to the quick-start protocol by the manufacturer. Briefly, 500 *µl* of QIAzol lysis reagent (#79306) was added to the isolates, mixed by pipetting, and incubated at room temperature (RT) for 5 min. After adding 100 *µl* chloroform, the tubes were capped securely and shaken vigorously for 15 *s*. The tubes were then incubated for 3 *min* at RT and centrifuged at 12,000 *g* for 15 *min* at 4 *°C*. The upper aqueous phase (∼300 *µl*), containing total RNA, was carefully transferred to a new 2 *ml* collection tube without transferring any interphase. 1.5 volumes of 100% ethanol were added (∼450 *µl*), and the samples were mixed thoroughly by pipetting. Up to 700 *µl* of sample was pipetted into Qiagen RNeasy MinElute spin column in a 2 *ml* collection tube, the tubes were capped and centrifuged at 8000 *g* for 15 *s* at RT. The flow-through was discarded. This step was repeated until all the sample passed through the spin column. 700 *µl* Buffer RWT (Qiagen #1067933) was added to each column, then the tubes were capped and centrifuged for 15 *s* at 8000 *g*, RT after which the flow-through was discarded. 500 *µl* Buffer RPE (Qiagen #1018013) was added to each column, the tubes were capped and centrifuged for 15 *s* at 8000 *g*, RT, then the flow-through was discarded. This step was repeated, but during the second time, the tubes were centrifuged for 2 *min* instead. The spin column was placed inside a new 2 *ml* RNAse-free collection tube and was centrifuged at full speed for 5 *min* at RT with the tube cap left open to dry the membrane. The collection tube containing the flow-through was discarded. Finally, the spin column was placed into a 1.5 *ml* RNAse-free collection tube, and 14 *µl* nuclease-free water was added directly to the centre of the column, then centrifuged at full speed at RT for 1 *min* to elute the RNA. The eluate was run through the column again to maximise the RNA recovery. The RNA was immediately placed on ice. RNA profile and concentration was then assessed by 2100 Bioanalyzer using RNA 6000 Pico kit (Agilent Technologies) according to manufacturer’s protocol.

### sEV surface protein profiling by flow cytometry

The MACSPlex Exosome kit (Miltenyi Biotec, Bergish Gladbach, Germany) utilizes beads functionalized with 37 types of antibodies to capture sEVs expressing the corresponding surface markers. The sEVs captured by the beads are then labelled with a cocktail of fluorescent labelled CD9, CD63 and CD81 antibodies which are detected by FCM. The expression of 37 surface markers (CD1c, CD2, CD3, CD4, CD8, CD9, CD11c, CD14, CD19, CD20, CD24, CD25, CD29, CD31, CD40, CD41b, CD42a, CD44, CD45, CD49e, CD56, CD62P, CD63, CD69, CD81, CD86, CD105, CD133, CD142, CD146, CD209, CD326, HLA-ABC, HLA-DRDPDQ, MCSP, ROR1, and SSEA-4) was evaluated by flow cytometry (FCM) with the short protocol specified by the manufacturer. Briefly, 5×10^8^ of particles/sample (calculated based on NTA measurement) or PBS (negative control) were diluted with MACSPlex buffer into 1.5 *ml* microcentrifuge tubes with a final volume of 120 *µl*. Each tube was incubated at room temperature for 1 *h* in the dark with 10 *µl* of MACSPlex exosome capture beads and 15 *µl* of the mix of APC-conjugated antibodies. Post-incubation, beads were washed twice with 500 *µl* of MPB at 3000 *g* for 5 *min*. Each time of washing, the supernatant was aspirated, leaving a residual volume of 150 *µl*. The prepared beads were then analysed by BD LSR Fortessa X-20 flow cytometer (BD Biosciences, San Jose, CA, USA). All mean fluorescence intensity (MFI) was background corrected according to the negative control.

For FCM comparison of sEV populations among plasma, SEC, SEC-DGUC and dUC, 5×10^8^ of particles/sample were mixed with 15 *µl* of MACSPlex exosome capture beads and incubated at room temperature for 1 hr in the dark. The beads were then washed thrice with 500 *µl* of MPB at 3000 *g* for 5 *min*. 15 µl of the mix of APC-conjugated antibodies was added to each sample and incubated at room temperature for 1 *h* in the dark. Post-incubation, beads were washed thrice with 500 *µl* of MPB at 3000 g for 5 *min*. Each time of washing, the supernatant was aspirated, leaving a residual volume of 150 *µl*. The prepared beads were then analysed by BD LSR Fortessa X-20 flow cytometer as described above.

### sEV protein profiling by liquid chromatography with tandem mass spectrometry

Mass spectrometry experiments were carried out at the proteomics core facility in the School of Biological Science at Nanyang Technological University. For SEC-DGUC-1, which was obtained from 5 *ml* of plasma, the first step was to remove the iodixanol through dialysis using Pur-A-Lyzer^TM^ Mini 6000 dialysis kit (PURN60030, Sigma-Aldrich, Co., STL, MO, USA). Then SEC-DGUC-1 was subjected to in-solution digestion prior to fractionation on an Xbridge™ C18 column (4.6 × 250 *mm*, Waters, Milford, MA, USA) and subsequent analysis by LC-MS/MS. SEC-PF was subjected to in-solution digestion and subsequent analysis by liquid chromatography with tandem mass spectrometry (LC-MS/MS).

For LC-MS/MS procedure, the peptides were separated and analyzed using a Dionex Ultimate 3000 RSLCnano system coupled to a Q Exactive instrument (Thermo Fisher Scientific, MA, USA). Separation was performed on a Dionex EASY-Spray 75 *μm* × 10 *cm* column packed with PepMap C18 3 *μm*, 100 *Å* (Thermo Fisher Scientific) using solvent A (0.1% formic acid) and solvent B (0.1% formic acid in 100% ACN) at a flow rate of 300 *nl/min* with a 60 *min* gradient. Peptides were then analyzed on a Q Exactive apparatus with an EASY nanospray source (Thermo Fisher Scientific) at an electrospray potential of 1.5 *kV*.

Raw data files were processed and searched using Proteome Discoverer 2.1 (Thermo Fisher Scientific). The Mascot algorithm was then used for data searching to identify proteins using the following parameters: missed cleavage of two; dynamic modifications were oxidation (+15.995 Da) (M) and Phosphorylation (+79.966 Da) (S, T, Y). The static modification was Carbamidomethyl (+57 Da) (C). Percolator was applied to filter out the false MS2 assignments at a strict false discovery rate of 1 % and relaxed false discovery rate of 5%. Only proteins identified with 2 or more peptides were included in the final list. The protein database used for protein identification was Uniprot Human.

## Author Contributions

Kong Fang and Megha Upadya planned, designed and performed the experiments. Kong Fang and Megha Upadya performed the sEV isolation and characterization. Kong Fang and Andrew See Weng Wong performed electron microscopy. Kong Fang, Megha Upadya and Rinkoo Dalan collected blood samples. Ming Dao and Kong Fang initiated, designed and conceptualized the study. Kong Fang, Megha Upadya and Ming Dao wrote the manuscript.

## Data Availability Statement

All data needed to evaluate the conclusions in the paper are present in the paper and/or the Supplementary Materials. Additional data related to this paper may be requested from the authors. Experimental procedures have been uploaded to EV-TRACK [31] with reference no. EV210379.

## Acknowledgement

This research has been funded through Nanyang Technological University Distinguished University Professorship awarded to S.S.. M.D. acknowledges partial support from National Institutes of Health under Grant No. R01HL154150. We appreciate the guidance from Prof. Bernhard Otto Boehm on the clinical aspects of sEV research. We thank Asst. Prof. Hou Han Wei for use of the Zetaview facility. We would like to express gratitude to Tan Tock Seng Hospital for providing the biobank samples.

## Supplementary Figure Data

**Supplementary Figure 1:**
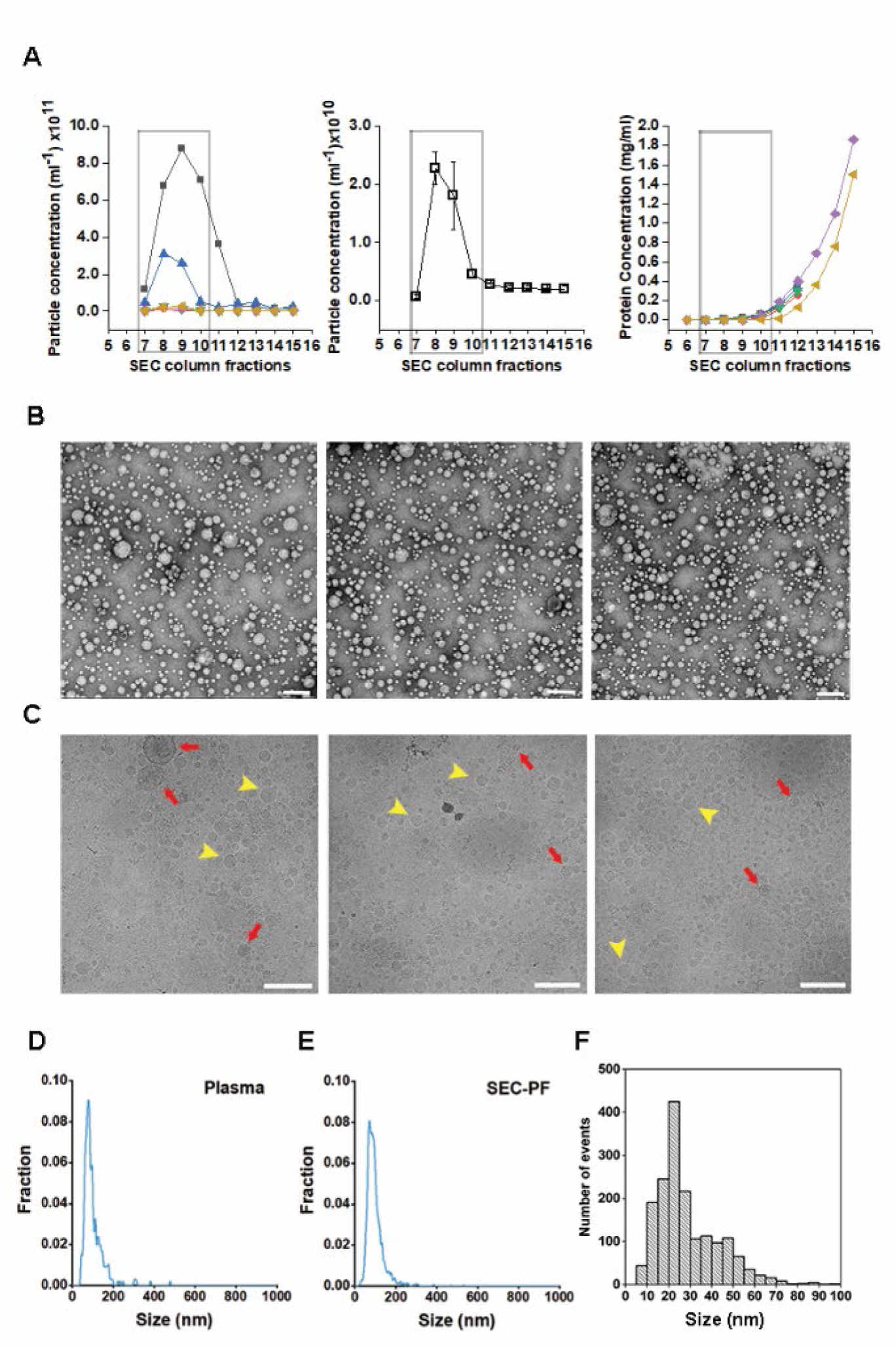
SEC elution profiles, EM images of the PF and particle size distributions. (A) SEC elution profiles according to particle concentration (by NTA) of 6 different plasma sources (left) and 4 repeats of the same plasma source (middle). Protein concentrations (by BCA, right) in corresponding to the elution profiles of the 6 different plasma sources shown on the left. 500 *μl* of plasma was loaded onto the SEC column and 10 fractions of 500 *μl* each were collected and analyzed. Fraction 7 to 10 were pooled to constitute the SEC-PF because of their high particle concentrations and low protein concentrations. (B) Representative TEM images of the PF. (C) Cryo-EM images of the SEC-PF. (D), (E) Typical particle size distributions of a plasma and SEC-PF measured by NTA. (F) A histogram of particle diameters obtained from the TEM images shown in (B).

**Supplementary Figure 2:**
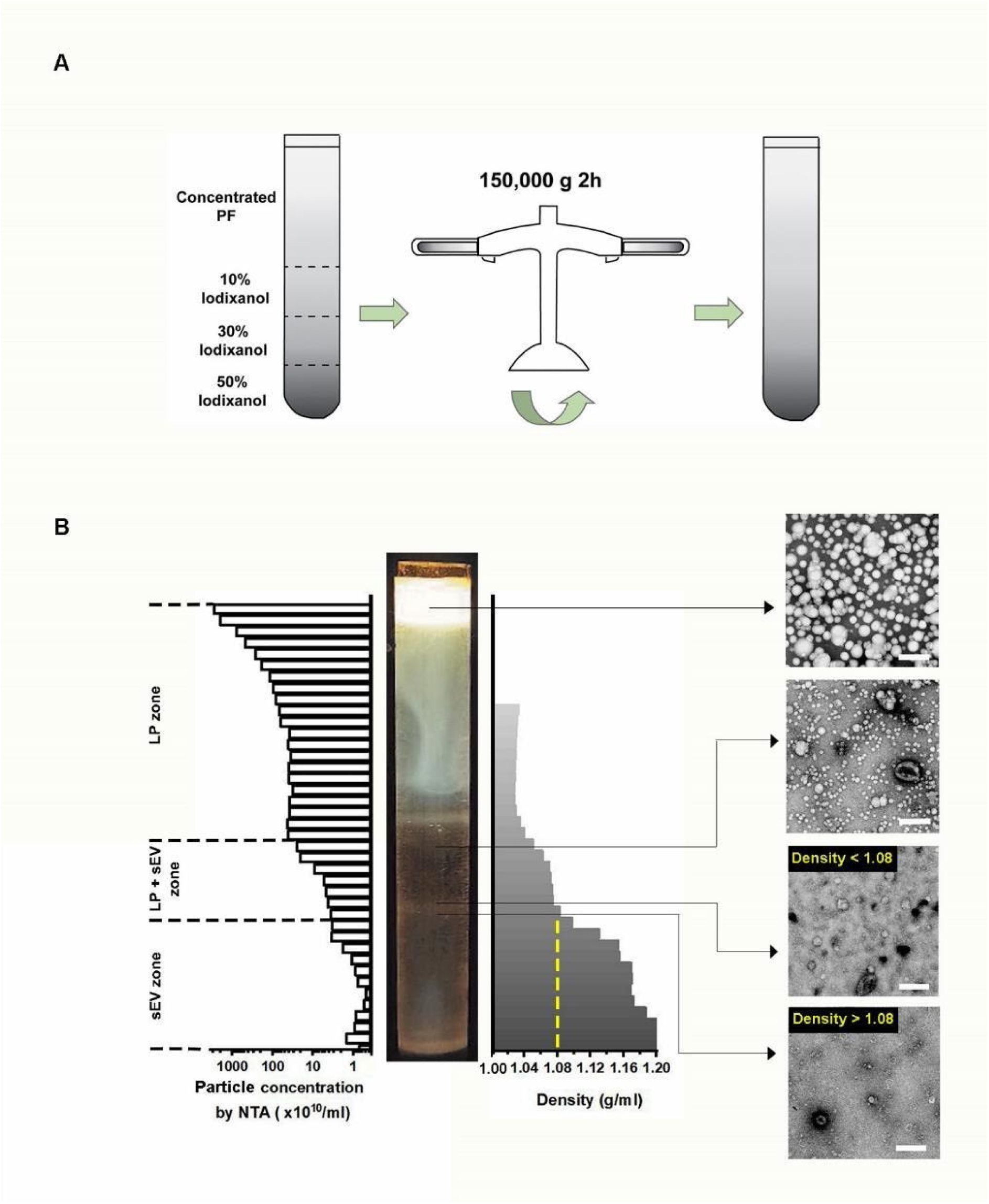
Distribution of sEVs and lipoproteins within the density gradient after DGUC. (A) SEC-PFs (6 *ml*) were placed on top of a density gradient cushion constructed with 2 *ml* 10%, 2 *ml* 30%, and 2 *ml* 50% iodixanol solutions and centrifuged at 150,000 x *g* for 2 *h* at 4 *°C*. (B) After DGUC, the tube was fractionated into 42 fractions, which were each examined for their densities, particle concentrations (by NTA) and presence of sEVs and lipoproteins (by TEM). The dominance of lipoproteins was evident in fractions of density <1.05 *g/ml,* which is designated as LP zone. sEVs started to appear when density exceeded 1.05 *g/ml* but a significant level of lipoproteins was present until the density of 1.08 *g/ml*. Therefore, the density region of 1.05∼1.08 *g/ml* was designated as LP+sEV zone. Beyond 1.08 *g/ml*, lipoproteins diminished drastically and the density region of >1.08 *g/ml* was designated as the sEV zone. All scale bars represent 200 *nm*.

**Supplementary Figure 3:**
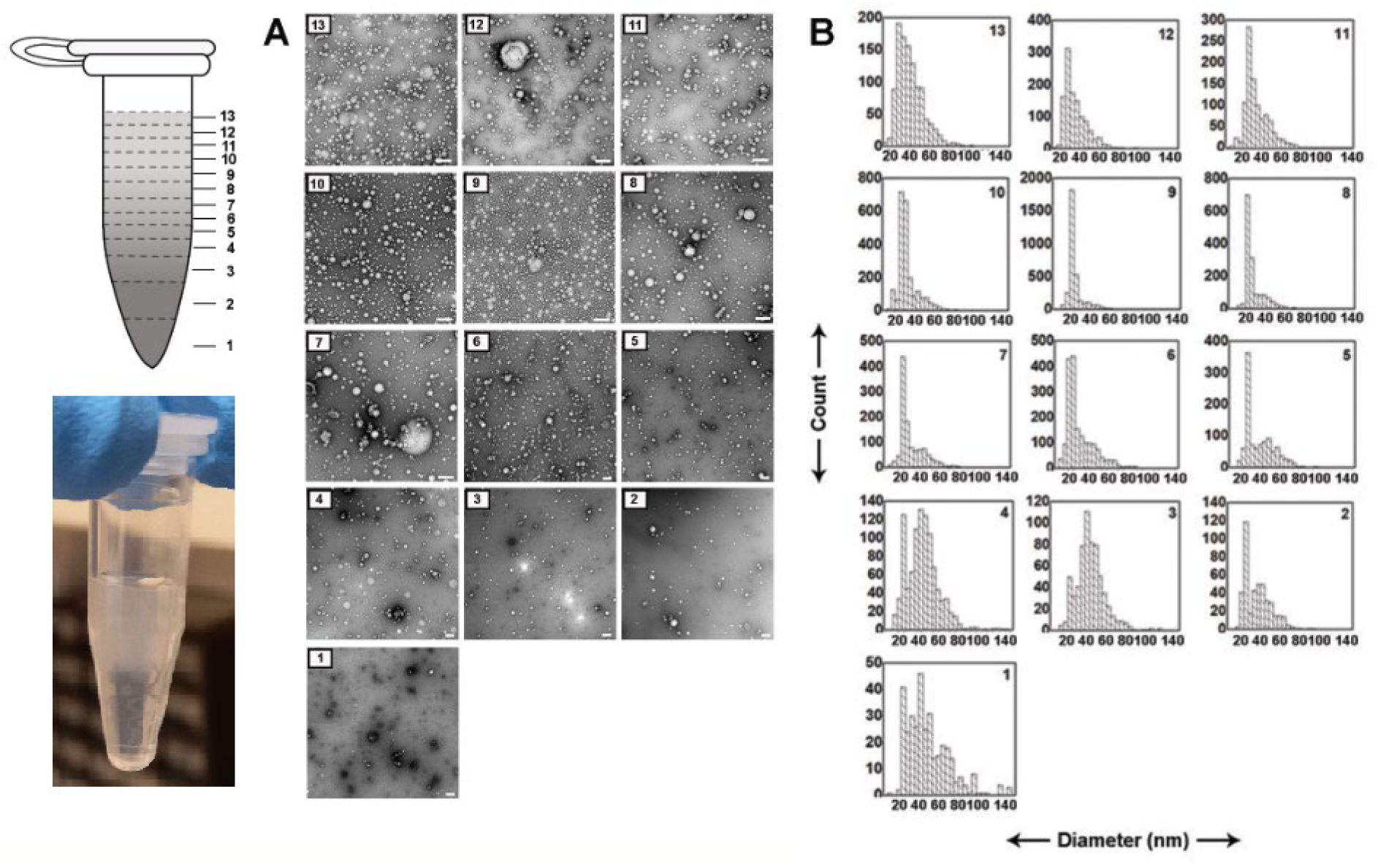
TEM images and size distributions of the 13 fractions collected from the 1.5 *ml* tube following SEC-DGUC. (A) Representative TEM images of each of the 13 fractions. SEC-DGUC-1 clearly showed minimum presence of lipoprotein (high contrast lighter color particles) compared to other fractions. Moreover, the presence of sEVs (low contrast and cup-shaped particles) is evident in SEC-DGUC-1. A typical image of the 1.5 *ml* tube after the DGUC step is shown at the left (bottom picture). The whitish layer at the top, gradually becoming clearer toward the bottom of the tube, corresponds well with the TEM observations. (B) Particle size distributions measured according to the TEM images of the 13 fractions.

**Supplementary Figure 4:**
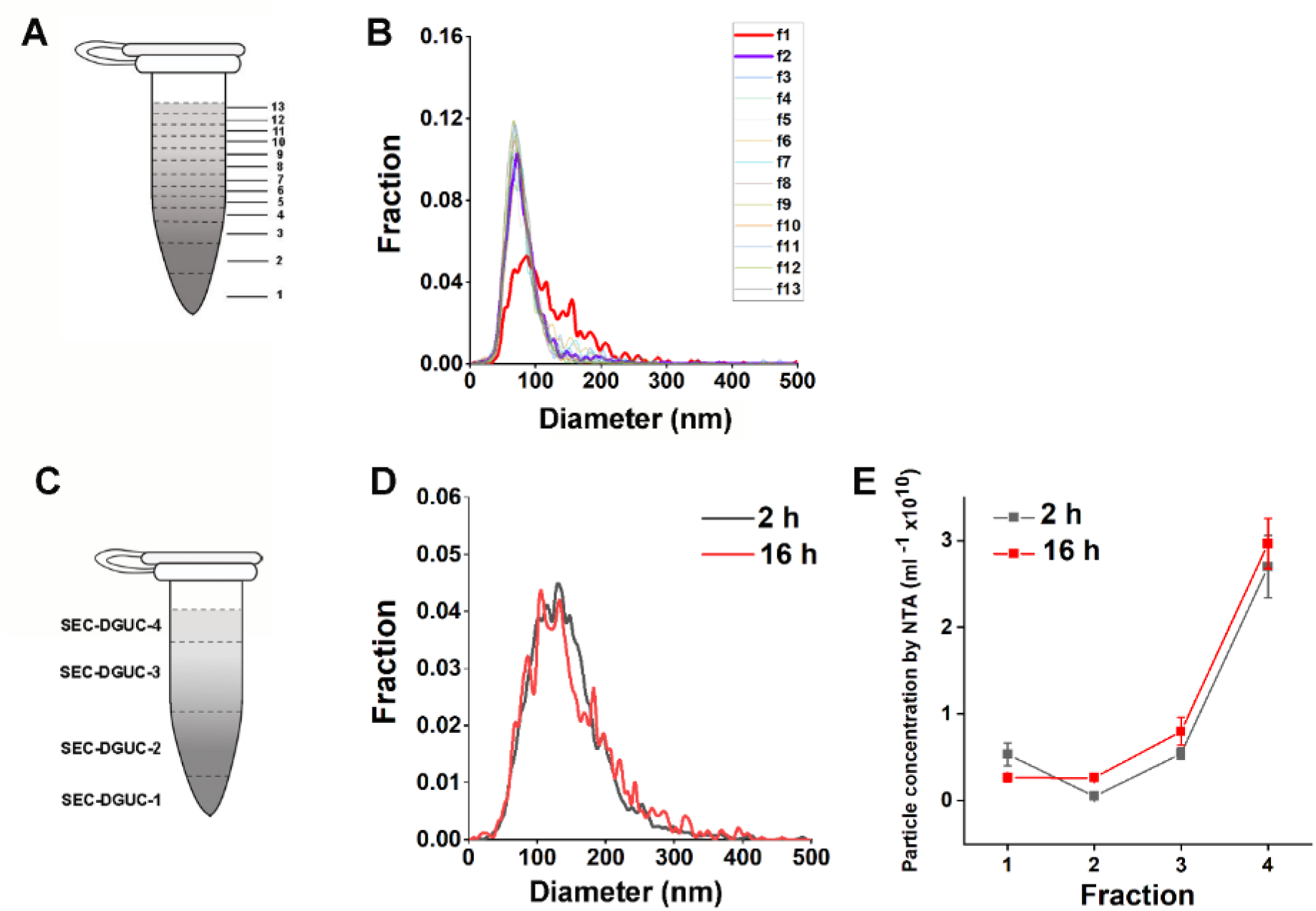
(A) The 1.5 *ml* tube was fractionated into 13 fractions. (B) Size distributions (by NTA) of the 13 fractions collected from the 1.5 *ml* tube following SEC-DGUC protocol. The particle size distribution in SEC-DGUC-1 was distinct from the rest of the fractions. Even in SEC-DGUC-2 (highlighted in purple), the size distribution follows fractions 3∼13, implying the dominance of lipoproteins in these fractions. (C) The 1.5 *ml* tube was fractionated into 4 fractions in order to examine if 2 *h* spinning time was sufficient to isolate sEV from lipoproteins in the 1.5 *ml* tube format DGUC. (D) The resulting size distributions of SEC-DGUC-1s (by NTA) were highly similar for 2 *h* and 16 *h* spinning time. (E) The particle concentrations of the four individual fractions along the 1.5 *ml* tube closely resembled each other for 2 *h* and 16 *h*, implying that there were no significant loss of sEV for 2 *h* spinning time in comparison to 16 *h*.

**Supplementary Figure 5:**
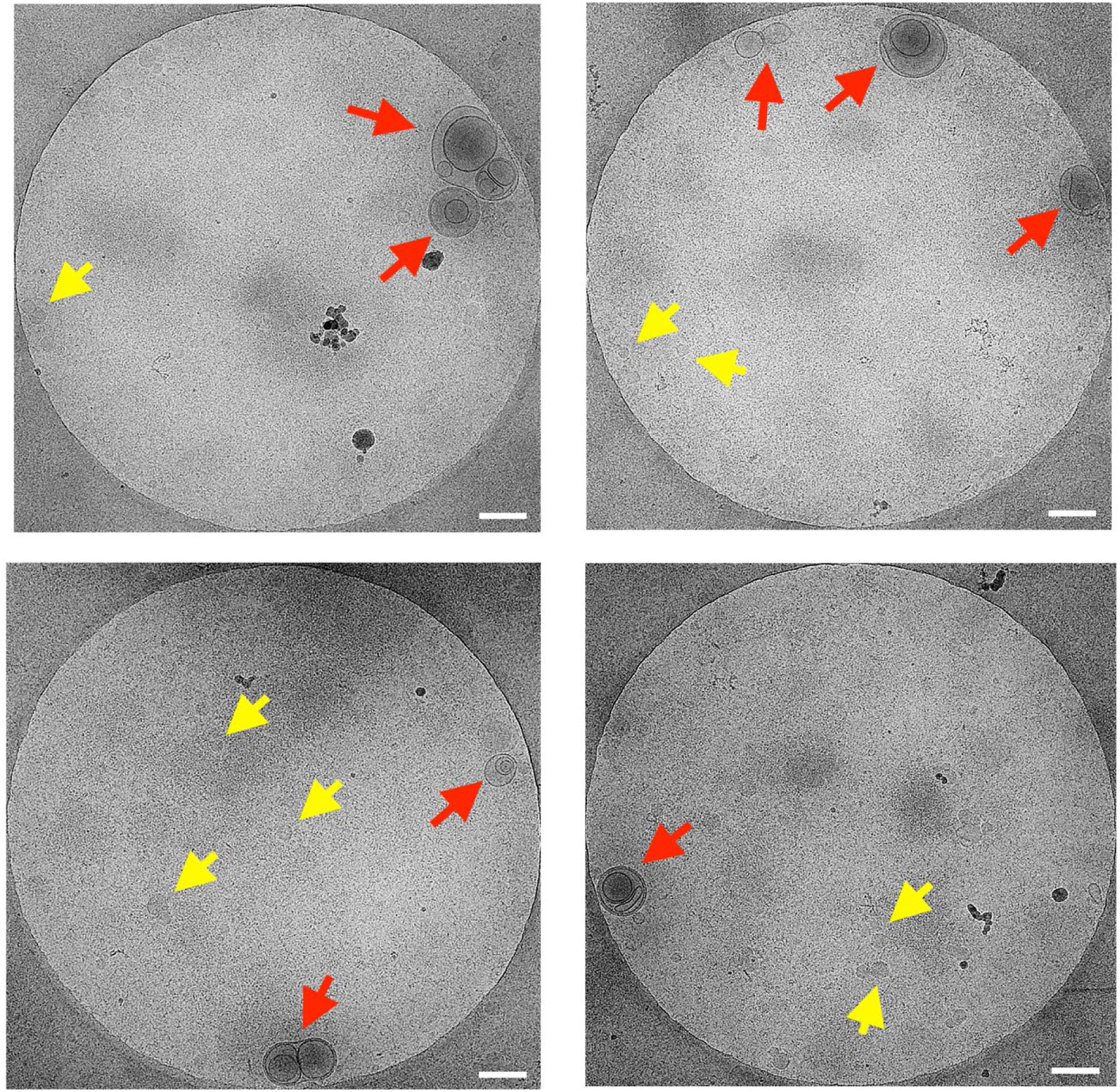
Additional Cryo-EM images of SEC-DGUC-1. Red arrows represent sEVs and yellow arrows represent typical lipoproteins. The SEC-DGUC-1 was obtained from non-fasting plasma collected in EDTA tubes. All scale bars represent 200 *nm*.

**Supplementary Figure 6:**
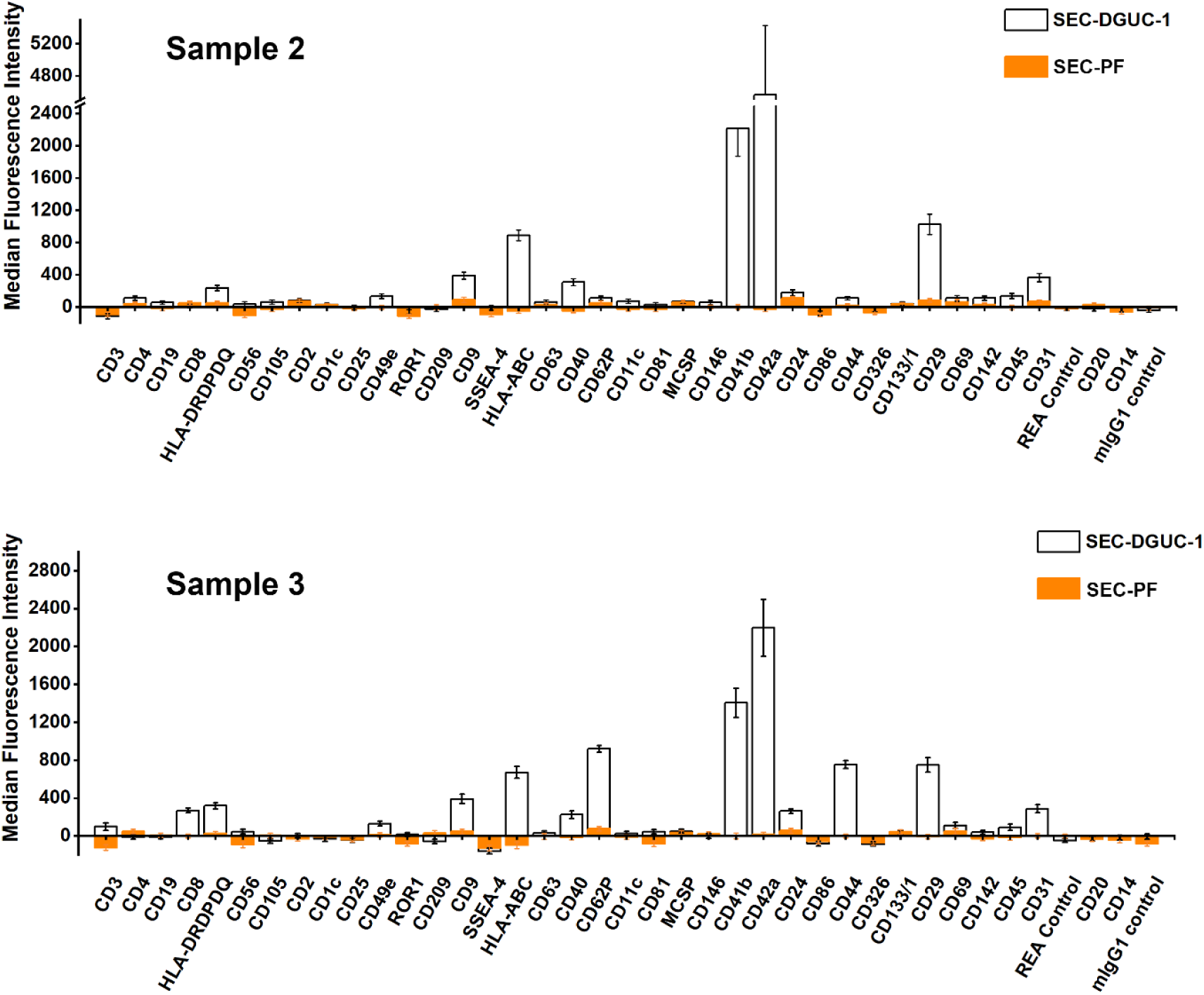
Flow cytometry data of SEC-PF vs. SEC-DGUC-1 obtained from two plasma sources using MACSPlex exosome kit. Data from two plasma sources demonstrated a similar trend of much stronger signals of SEC-DGUC-1 compared to SEC-PF in flow cytometry using MACSPlex exosome kit. Data are Median ± SEM.

**Supplementary Figure 7:**
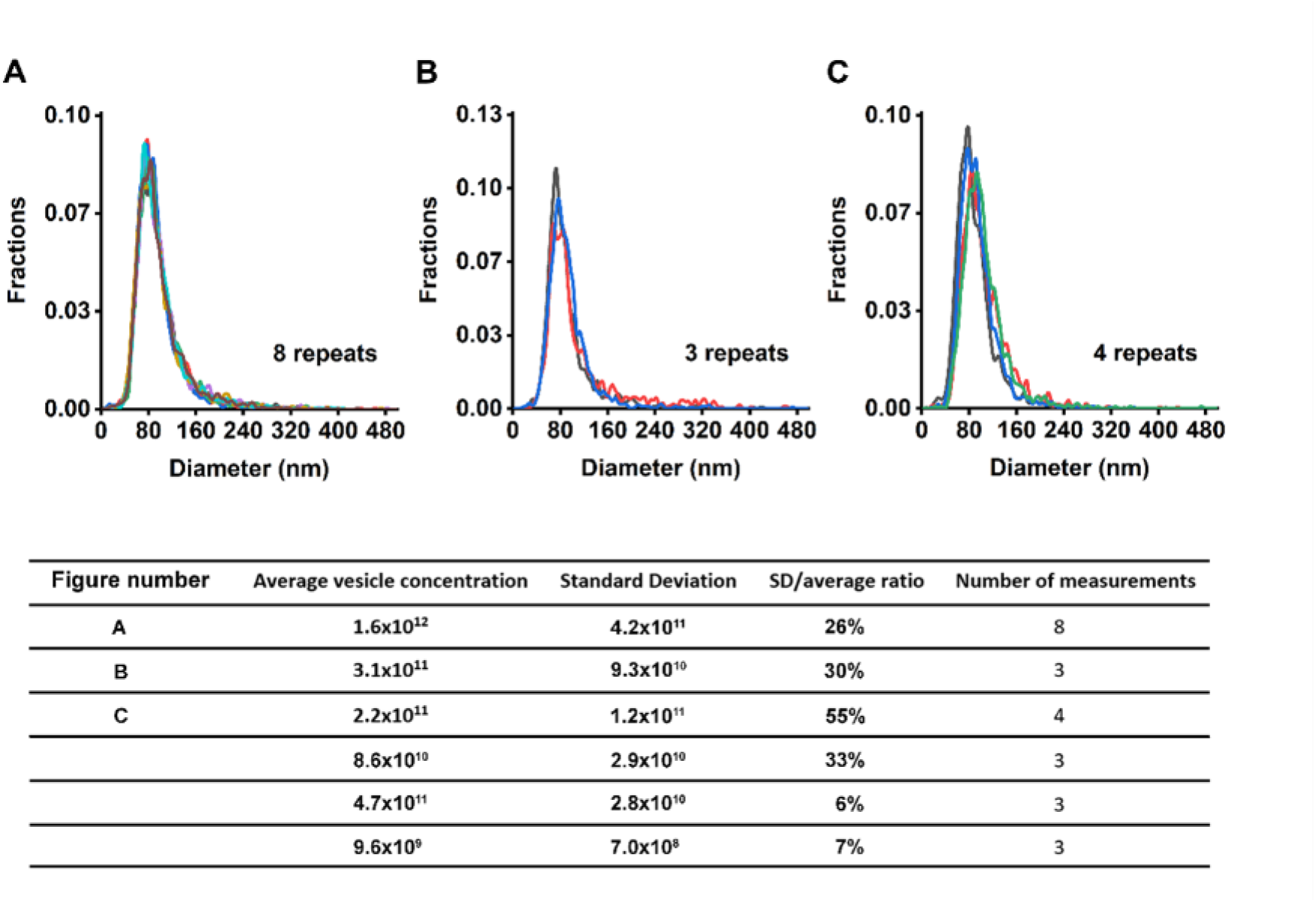
Repeatability of SEC. (A-C) Particles size distributions of SEC-PF measured by NTA. (A), (B) and (C) correspond to the first, second and third rows of the table, respectively. The highly overlapped size distributions allude to the consistency in the particles eluted from SEC columns, even though the number of particles eluted varied up to 55%. The table lists six experiments with various numbers of repeats of SEC. The average particle concentrations, standard deviation and CV (SD/average ratio) are listed together with the number of measurements made.

**Supplementary Figure 8:**
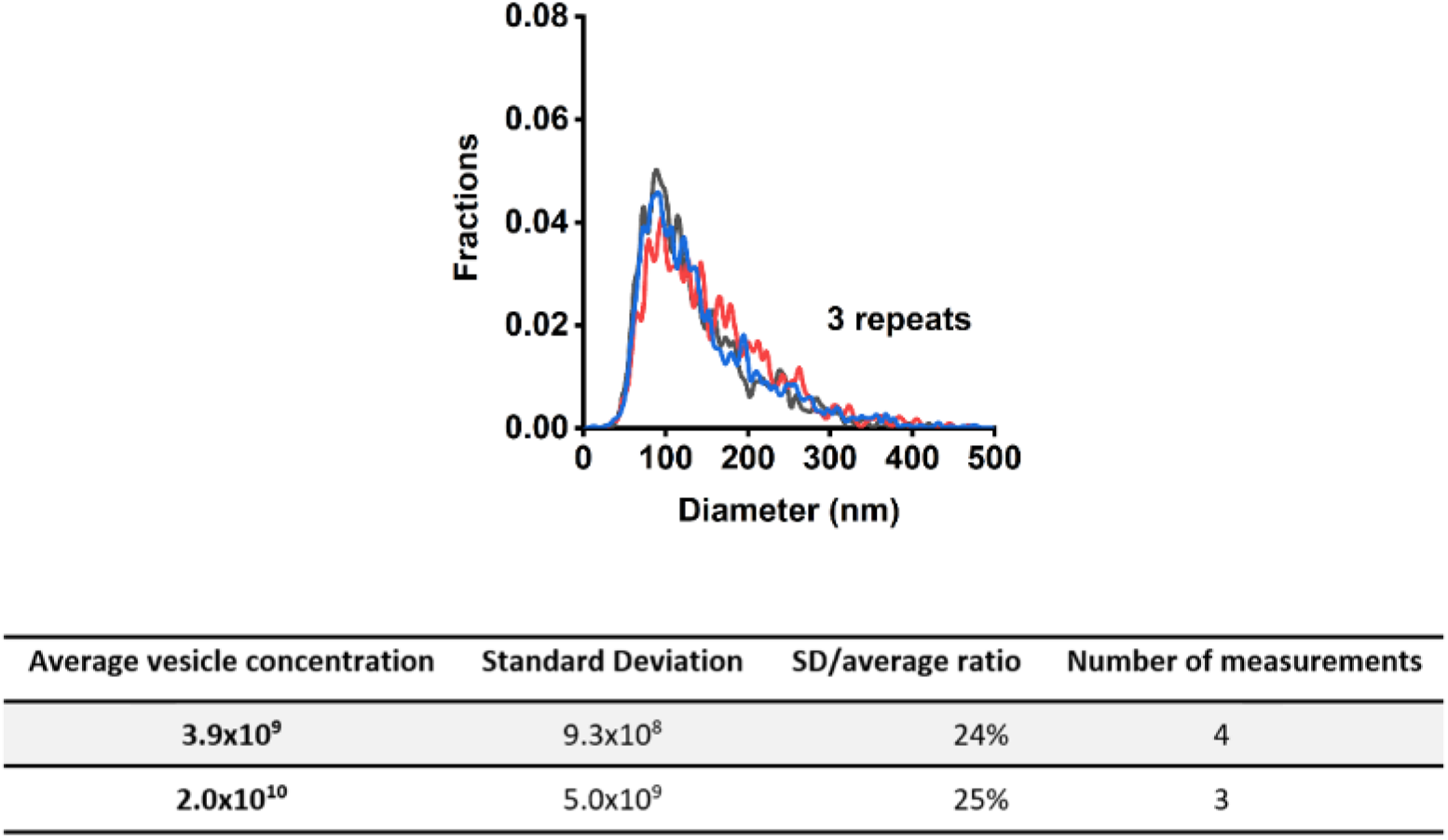
Repeatability of SEC-DGUC protocol. NTA measurements of two experiments to test repeatability after SEC-DGUC. The figure displays the size distributions measured by NTA of the experiment listed in the second row of the table. The size distributions of the experiment listed in the first row of the table were presented in main Figure 9. The table lists the average particle concentrations of SEC-DGUC-1 and the standard deviation, CV (SD/average ratio) together with the number of measurements made.

**Supplementary Figure 9:**
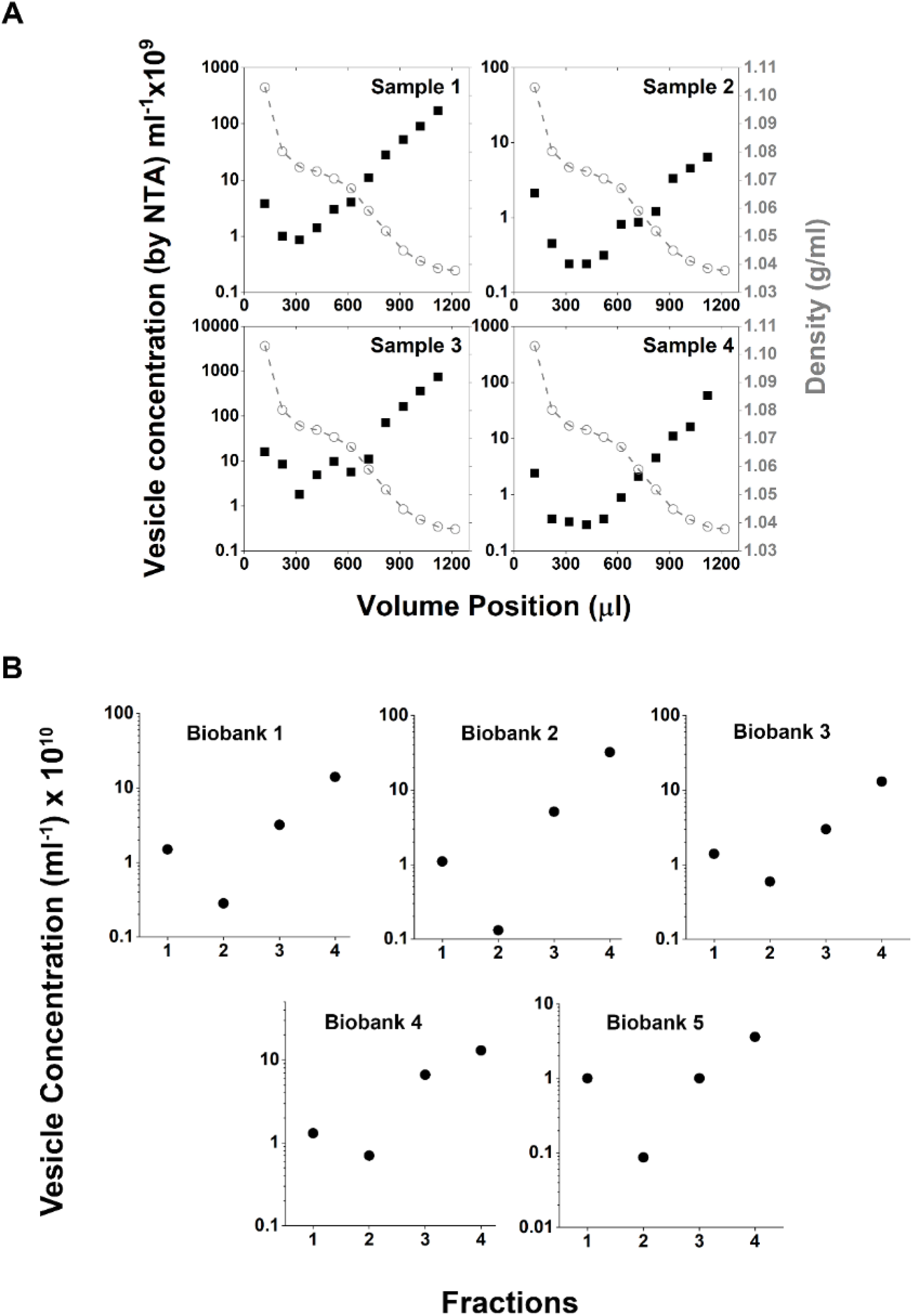
Reliability of particle concentration profiles along the 1.5 *ml* tube (by NTA). (A) Detailed particle concentration profiles (13 fractions) of four different samples along with the calibrated density profile shown in Figure 2. (B) Five particle concentration profiles (4 fractions shown in Figure 9B) in SEC-DGUC-1 corresponding to the biobank plasma samples (EDTA tubes) shown in Figure 11. Note that the data from (A) and (B) are from different plasma sources.

**Supplementary Figure 10:**
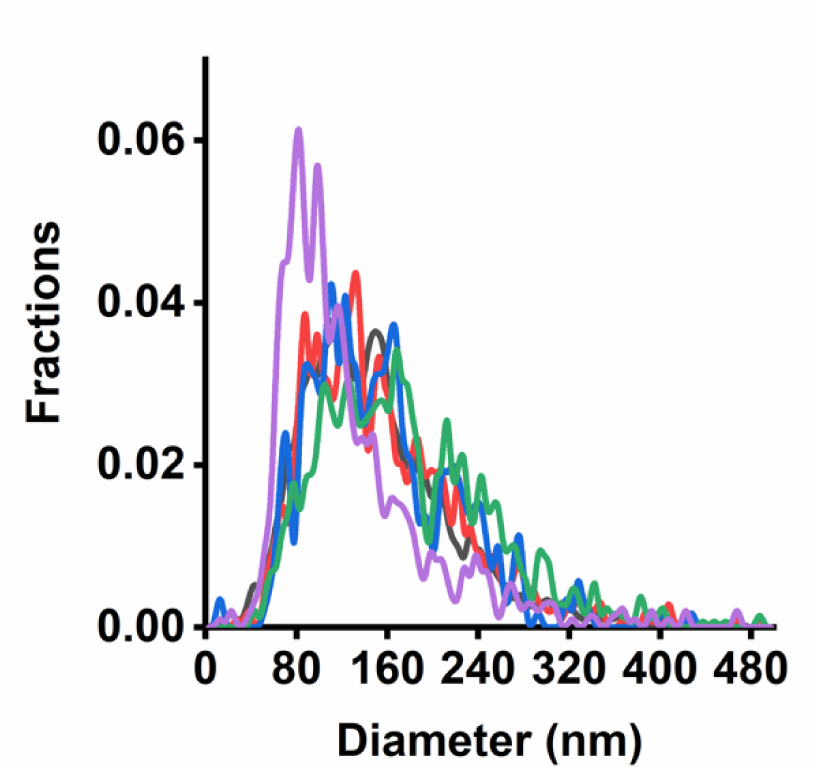
Size distributions (by NTA) of SEC-DGUC-1 obtained from 5 fasting plasma corresponding to Figure 11. The particle size distributions did not overlap, reflecting the variation of sEV population among different plasma sources.

